# Proteome-Wide Photo-Crosslinking Enables Residue-Level Visualization of Protein Interaction Networks *in vivo*

**DOI:** 10.1101/2022.09.20.508727

**Authors:** Anneliese M. Faustino, Piyoosh Sharma, Divya Yadav, Stephen D. Fried

**Author notes:** Contributed equally to this work (Anneliese M. Faustino, Piyoosh Sharma).

## Abstract

Crosslinking mass spectrometry (XL-MS) is emerging as a unique method at the crossroads of structural and cellular biology, uniquely capable of identifying protein-protein interactions with residue-level resolution and on the proteome-wide scale. With the development of crosslinkers that can form linkages inside cells and easily cleave during fragmentation on the mass spectrometer (MS-cleavable crosslinks), it has become increasingly facile to identify contacts between any two proteins in complex samples, including in live cells or tissues. Photo-crosslinkers possess the advantages of high temporal resolution and high reactivity, thereby engaging all residue-types (rather than just lysine); nevertheless, photo-crosslinkers have not enjoyed widespread use, and have yet to be employed for proteome-wide studies, because their products are challenging to identify, and an MS-cleavable photo-crosslinker has not yet been reported. Here, we demonstrate the synthesis and application of two heterobifunctional photo-crosslinkers that feature diazirines and *N*-hydroxy-succinimidyl carbamate groups, the latter of which unveil MS-cleavable linkage upon acyl transfer to protein targets. Moreover, these crosslinkers demonstrate high water-solubility and cell-permeability. Using these compounds, we demonstrate the feasibility of proteome-wide photo-crosslinking mass spectrometry (photo-XL-MS), both in extracts and *in cellulo*. These studies provide a partial interaction map of the *E. coli* cytosol with residue-level resolution. We find that photo-XL-MS has a propensity to capture protein-protein interactions, particularly involving low-abundance uncharacterized proteins, suggesting it could be a powerful tool to shed light on the “darker” corners of the proteome. Overall, we describe methods that enable the detection of protein quinary interaction networks in their native environment at residue-level resolution proteome-wide, and we expect they will prove useful toward the effort to explore the molecular sociology of the cell.

**TOC graphic:** 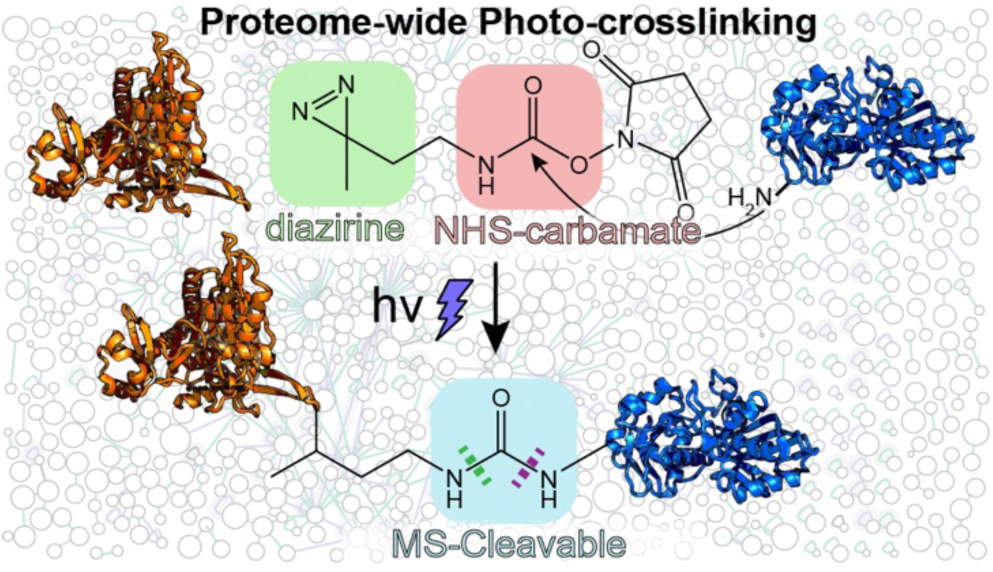

## INTRODUCTION

With recent technical and technological advances, crosslinking mass spectrometry (XL-MS) is emerging as a powerful structural proteomics technique – uniquely capable of elucidating protein-protein interaction (PPI) networks at residue-level resolution and on the proteome-wide scale.^1–4^ Specifically, improvements in high-resolution mass analyzers, chemical crosslinkers, and analysis software have joined forces to move XL-MS from a structural biology method (applied to purified protein complexes) to an “omics” technique that bridges between cellular and structural biology. For example, in a recent report, O’Reilly and co-workers combined in-cell XL-MS with cryo-electron tomography to develop a structural model of the RNA polymerase-ribosome complex performing coupled transcription-translation *in situ*.^5^

Early crosslinkers applied in XL-MS studies, such as disuccinimidyl suberate (DSS) and bis(sulfosuccinimidyl)suberate (BS3), are homobifunctional compounds consisting of two NHS-esters connected by a hydrocarbon linker.^6–11^ Following chemical crosslinking and trypsinolysis, these compounds generate pairs of peptides (called crosslinked peptides) irreversibly adjoined between two nucleophilic residues (typically lysines). Developed in the late 2000s, these crosslinkers proved useful for obtaining high coverage on single proteins; however, they were challenging to apply to complex samples, such as lysates, cells, or tissues. This is because the complexity of the search space associated with possible crosslinked peptides scales as the number of linear peptides squared (known as the *N*^2^ problem),^12^ which complicates spectral identification and raises the chances of false discovery.

Proteome-wide XL-MS studies emerged with the onset of a newer generation of crosslinkers,^13, 14^ such as disuccinimidyl sulfoxide (DSSO) and disuccinimidyl dibutyric urea (DSBU, shown in Figure 1A), which can be cleaved in the gas phase during collision-induced dissociation (CID) or higher-energy collision-induced dissociation (HCD). These crosslinkers feature a functional group (cyan boxes in Figure 1) which is readily broken, splitting the crosslinked peptide into two modified linear peptides. In a symmetric crosslinker, fissure of the functional group can occur in one of two ways, which further facilitates the identification of spectra of crosslinked peptides by generating characteristic twin peaks (Figure 1C). These features collectively enable MS-cleavable crosslinkers to mitigate the *N*^2^ problem by reducing the database size against which spectra are searched and allowing complex samples to be interrogated by XL-MS. While numerous functional groups have been reported to be MS-cleavable,^14^ sulfoxides and urea are among the most frequently employed.^15, 16^ With these chemical tools, several proteome-wide crosslinking studies have been reported, such as on *Drosophila* embryonic extracts,^16^ *Mycoplasma pneumoniae* cells,^5, 17^ murine mitochondria,^18^ murine heart tissues,^19^ human adenocarcinoma extracts,^20^ and in HEK-293 cells.^21^

**Figure 1.**
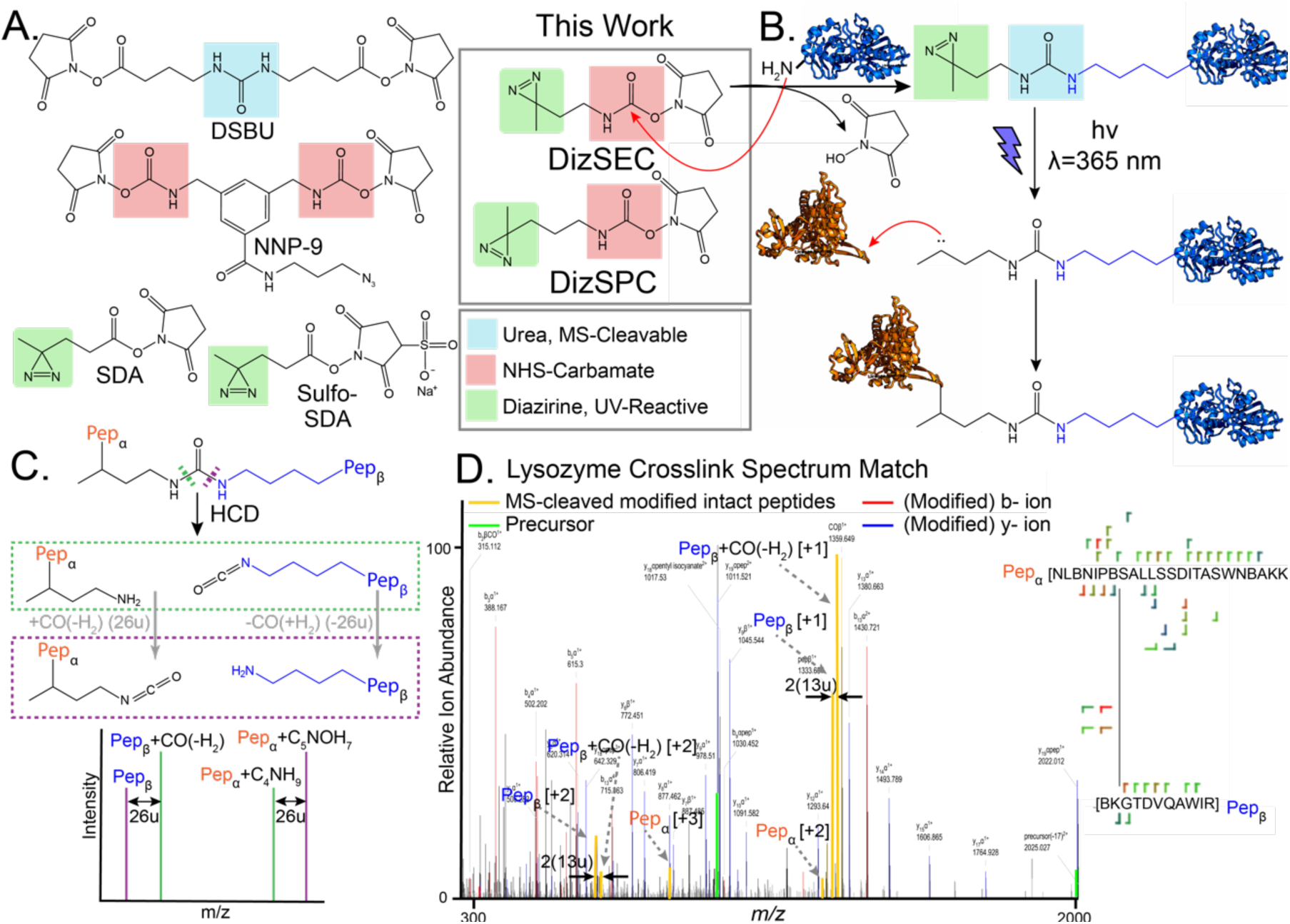
Structure and Properties of MS-cleavable Heterobifunctional Photo-crosslinkers. (**A**) Structures of DSBU, a urea-based (teal box) cleavable crosslinker; NNP-9, an NHS-carbamate (red box) containing-crosslinker; and SDA, a heterobifunctional diazirine (green box) crosslinker. Shown on the right are the heterobifunctional NHS-carbamate photo-crosslinkers, DizSEC and DizSPC, reported in this study. (**B**) Reaction of the NHS-carbamate functional group in DizSEC or DizSPC with lysine or N-terminal amines generates ureas. Following UV irradiation, the diazirine forms a carbene, which rapidly reacts with proximal proteins to form covalent crosslinks. (**C**) Ureas can be cleaved under HCD in two different but symmetrical ways (green and purple dashed boxes) to generate amine and isocyanate fragments on either peptide product (orange α peptide and blue β peptide). These crosslinked peptide fragments form characteristic twins of doublets in the MS spectra. (**D**) A representative crosslinked spectrum match (XSM) generated by the treatment of lysozyme with DizSEC. The spectrum contains three of the four ions (yellow) that correspond to the cleaved, modified, but otherwise unfragmented parent peptides.

Photo-crosslinkers (also called photo-affinity labels (PALs)) possess chemically inert photo-reactive functional groups, which upon UV irradiation, generate highly reactive moieties that rapidly and covalently capture nearby proteins.^22, 23^ Though azides, benzophenones and diazirines are the most utilized PALs in biological systems, recent studies have reported aryl-tetrazoles and o-nitrobenzyl alcohols as PALs to capture protein-protein interactions.^24, 25^ Diazirines are “zero-length” crosslinkers that are stable in aqueous media at room temperature, are synthetically tractable in a broad range of architectures, and offer predictable photo-crosslinking products; hence, they have emerged as a popular “universal” photo-affinity label.^26–28^ Additionally, diazirines are photo-activated at relatively benign UV wavelengths (∼365 nm), avoiding protein damage that can occur at shorter wavelengths (< 300 nm).^29^ Upon photoactivation, diazirines expel N_2_ and form reactive carbenes that can insert across a diverse array of chemical bonds, including N-H, C-H, O-H, and S-H.^30^ Aliphatic carbenes evolved from diazirines have short lifetimes (ns–µs) and are reactive toward any amino acid in proximity. Hence this combination of properties enables diazirene-based photo-crosslinkers to capture rapid or transient interactions and significantly expands “crosslinkable” chemical space, given that most chemical crosslinkers attach to a restricted profile of residue-types (primarily Lys or Cys).^23^

Heterobifunctional photo-crosslinkers such as succinimidyl-4,4’-azipentanoate (SDA) and sulfo-SDA (Figure 1A) contain an amine-reactive NHS-ester and a diazirine (green box) separated by an alkyl chain. This family of crosslinkers was initially introduced to assist investigations of PPIs by covalently coupling interacting proteins to a target protein of interest.^31–34^ They utilize a “plant-and-cast” strategy whereby the amine-reactive NHS-ester^35^ plants the compound onto a target protein, casting the photo-crosslinking moiety to react with proximal residues upon subsequent UV irradiation. The crosslinks generated by heterobifunctional photo-crosslinkers can also be sequenced with mass spectrometry; however, identifying photo-crosslinked peptides is technically challenging because: (i) their higher reactivity results frequently in quenching and consequently lower crosslinking yields; and (ii) they can react with all residue-types through a variety of linkages.^36, 37^ Hence, photo-crosslinkers have found some application in structural studies to generate interaction maps spanning diverse residue types (yielding orthogonal and complementary information to what is obtained with conventional chemical crosslinkers);^38, 39^ however, the challenges associated with photo-crosslinking have resulted in limited applicability to proteome-wide experiments. Indeed, SDA and sulfo-SDA lack MS-cleavable functional groups, which precludes confident identification of crosslinked peptides from complex search spaces.

To our knowledge, a heterobifunctional photo-crosslinker that produces an MS-cleavable crosslink has not been previously described. We reasoned that this combination of characteristics could enable the rapid capture of transient protein-protein interaction networks and then facilitate their identification from complex samples. We were inspired by the compound NNP-9 (Figure 1A), a trifunctional chemical crosslinker that features an NHS-carbamate group (red box).^40^ Following this concept, we designed two heterobifunctional crosslinkers like SDA, but in which the NHS-ester moiety is replaced with an NHS-carbamate. Following a nucleophilic attack by lysine, NHS-carbamates yield urea functional groups at the initial attachment site. This group would possess MS-cleavability, which could facilitate spectral search (Figure 1B), though this latent capacity of NHS-carbamates has not been previously exploited.

Here we report the synthesis and evaluation of DizSEC and DizSPC, photo-crosslinkers featuring NHS-carbamates and aliphatic diazirines with ethyl and propyl spacer arms, respectively, separating the chemical and photo crosslinking moieties. In the following, we demonstrated that DizSEC and DizSPC could generate readily identifiable crosslinks in pilot studies on purified proteins. We then provide evidence that MS-cleavability, combined with an optimized workflow, enables high-volume identification of crosslinks in whole *E. coli* extracts. Finally, we demonstrate that DizSEC and DizSPC are cell-permeable and develop a method to perform rapid *in vivo* photo-crosslinking of whole *E. coli* cells. The high number of crosslinks identified generates a partial interaction map of the *E. coli* cytosol with residue-level resolution, capturing many previously-unknown – and likely transient – protein-protein interactions that feature uncharacterized proteins. Therefore, the compounds and methods described in the following provide a foundation to perform proteome-wide photo-crosslinking mass spectrometry, which we hope will become a valuable tool to elucidate the molecular sociology of the cell with high temporal, spatial, and compositional resolution.^41, 42^

## EXPERIMENTAL METHODS

### Synthetic Methods

Heterobifunctional photo-crosslinkers DizSEC and DizSPC were synthesized in a seven-step protocol from commercial reagents with yields of 86.7% (111 mg) and 85.9% (116 mg), respectively. Detailed synthetic procedures and characterization are reported in Supporting Information (Scheme S1, Figures S1-S8).

### Crosslinking Lysozyme with DizSEC and DizSPC

Hen egg-white lysozyme (10 µM, Sigma-Aldrich) was prepared in reaction buffer (50 mM Na-HEPES pH 8.0, 100 mM NaCl) at a 50 µL scale to which 0.5 µL of freshly prepared photo-crosslinkers DizSEC, DizSPC, or sulfo-SDA (as positive control) were added as stocks in DMSO (100 mM or 1 M stock concentration) to a final concentration of 1 mM or 10 mM. Some samples received 4-dimethylaminopyridine (DMAP) to a final concentration of 0.1 mM or 1 mM.^43, 44^ Samples were gently mixed by inversion, then incubated at room temperature on an end-over-end rotator for 2 h (10 rpm). Chemical crosslinking reactions were quenched by the addition of Tris pH 8.0 (100 mM, final concentration) followed by incubation for 15 min at room temperature. Samples were transferred to a 1.0 mm path length quartz cuvette and photolyzed for 5.0 s with a 4.0 cm^2^ Hönle LED Spot W UV (365 nm)-LED array rated with a 14 W/cm^2^ max output (20% or 80% intensity). The distance between the cuvette and the LED head was minimized (ca. 1–2 mm) by affixing the cuvette with tape to the array. Tables S1 and S2 (Supporting Information) catalog various experimental crosslinking conditions used for the lysozyme samples.

### Proteome-Wide Crosslinking of *E. Coli* Extracts

*E. coli* cells (K12 subst. MG1655) were grown overnight in LB (5 mL) at 37 °C, 220 rpm. Overnight cultures were diluted into 50 mL LB with an initial OD_600_ of 0.05 until a final OD_600_ of 1.0 was reached. The cultures were pelleted in a 5910R swing-bucket benchtop centrifuge (Eppendorf) at 4000 *g* for 10 mins (4 °C), and the supernatant was discarded. The cell pellet was vigorously resuspended in 1 mL of HEPES reaction buffer and then lysed by sonication on water-ice with an FB120 probe sonicator (Fisher Scientific) at 45% amplitude for 10 min in 10 s/10 s on-off cycles. *E. coli* lysates were centrifuged at 14,000 *g* for 10 minutes on a 5425R benchtop centrifuge (Eppendorf) to clarify the samples. BCA assay (Pierce) was performed following a manufacturer’s protocol to determine the protein concentration. Samples were normalized to 1.0 mg/mL protein with reaction buffer. The crosslinking reaction was performed on a 1 mL scale of the normalized extracts. DizSEC or DizSPC (10 mM final concentration) were added from 1 M stocks in DMSO. Some samples were also supplied with 1 mM DMAP (final concentration) as a crosslinking catalyst. Samples were gently mixed by inversion, then incubated at room temperature on an end-over-end rotator for 2 h (10 rpm). Chemical crosslinking reactions were quenched by the addition of Tris pH 8.0 (100 mM, final concentration) followed by incubation for 15 min at room temperature. Samples were transferred to a 1.0 mm path length quartz cuvette (Starna Cells, PN: 21-Q-1) and photolyzed for 5.0 s with a 4.0 cm^2^ Hönle LED Spot W UV (365 nm)-LED array rated with a 14 W/cm^2^ max output (20% or 80% intensity). The UV-LED array was connected to a 200 W LED PowerDrive (Hönle).

### In-Cell Photo-Crosslinking of *E. coli*

Overnight and day cultures were prepared identically to that described above. 50 mL LB cultures were pelleted in a 5910R swing-bucket benchtop centrifuge (Eppendorf) at 4000 *g* for 10 mins (4 °C), and the supernatant was discarded. Cells were resuspended and washed in 20 mL of HEPES reaction buffer four times. After the final centrifugation, cells were resuspended in 1 mL HEPES reaction buffer, and the NHS-carbamate coupling reactions were performed by the addition of DizSEC or DizSPC (10 mM final concentration). Cell suspensions were gently mixed by inversion, then incubated at room temperature on an end-over-end rotator for 2 h (10 rpm). Cell suspensions were diluted 10-fold by the addition of 9 mL HEPES reaction buffer before photolysis to prevent excessive scattering.

The photolysis apparatus consisted of the LED Spot W UV-LED array with a 1 mm pathlength quartz flow-cell (Starna Cells, PN: 45-Q-1) affixed directly to the LED array surface with tape. The flow cell was connected to a Masterflex L/S Model 77200-50 peristaltic pump via 1.6 mm ID C-Flex biocompatible tubing (Masterflex). The tubing and flow cell were primed with water at max speed to remove bubbles. The cell suspension was pumped through the flow cell at 50 rpm (approximately 1 s exposure time), collected, quenched by incubation with Tris pH 8 (100 mM, final concentration) for 15 minutes at room temperature, and centrifuged at 4000 *g* for 10 min (cf. Figure 4A). The crosslinked cells were resuspended in 1 mL HEPES reaction buffer, then lysed by sonication and normalized to 1.0 mg/mL protein using the BCA assay as described above.

### MS Sample Preparation

Crosslinked protein mixtures (typically 1 mL of 1.0 mg/mL) were denatured by addition of urea (1.92 g), ammonium bicarbonate (Ambic, 15.8 mg) and diluted to a final volume of 4 mL by addition of Millipore water to generate samples that are 0.25 mg/mL protein, 8 M urea, and 50 mM Ambic (pH 8). These samples were reduced by addition of dithiothreitol (DTT) to 10 mM final concentration and incubation at 37 °C with agitation (700 rpm) in a benchtop ThermoMixer (Eppendorf) for 45 minutes. Samples were then alkylated by addition of iodoacetamide (IAA) to 40 mM final concentration and incubation at room temperature in the dark for 30 min. Samples were either: (i) diluted with 3 volumes of 50 mM Ambic and digested with trypsin (1:50 enzyme:protein w/w ratio, Pierce) overnight (ca. 16 h) at 25 °C, 700 RPM in a ThermoMixer; or (ii) digested serially – first with Lys-C (1:100 enzyme:protein w/w ratio, Pierce) for 2 h at 37 °C, then diluted with 3 volumes of 50 mM Ambic followed by second digestion with trypsin overnight (ca. 16 h). Desalting of protein digests was performed using Sep-Pak C18 cartridges (Waters).

Lysozyme samples were desalted with C18 cartridges (Waters Corporation). First, digests were acidified with HPLC-grade trifluoroacetic acid (TFA, Alfa Aesar) to a final concentration of 1% v/v and diluted to 1 mL with 0.5% TFA in Optima LC/MS grade water (Fisher chemical, Buffer A). Cartridges were placed onto a vacuum manifold, conditioned with 2 x 1 mL 0.5% TFA in 80% Optima LC/MS grade acetonitrile (Fisher chemical, Buffer B), and equilibrated with 4 x 1 mL Buffer A. Acidified digests were loaded onto the cartridges under a reduced vacuum (such that it took approximately 5 min for 1 mL to passage through the sorbent) and then washed with 4 x 1 mL Buffer A. The cartridges were placed on top of 15 mL conical tubes, and 1 mL of Buffer B was pipetted onto the resin. The cartridges were gently eluted under centrifugal force in swing-buckets spun at 350 rpm in a 5910R centrifuge (Eppendorf) for 3 min and then reduced to dryness in a Vacufuge centrifugal concentrator (Eppendorf).

*E. coli* extracts and in-cell crosslinking samples were desalted analogously, except using high-capacity C18 cartridges (Waters Corporation). Samples were acidified as described above. High-capacity cartridges were placed onto a vacuum manifold, conditioned with 2 x 10 mL Buffer B, and equilibrated with 4 x 10 mL Buffer A. Acidified digests were loaded onto the cartridges under a reduced vacuum (such that it took approximately 5 min for the ca. 12 mL to passage through the sorbent) and then washed with 4 x 10 mL Buffer A. The cartridges were placed on top of 50 mL conical tubes, and 5 mL of Buffer B was pipetted onto the resin. The cartridges were gently eluted under centrifugal force in swing-buckets spun at 350 rpm on a 5910R centrifuge for 5 min and then reduced to dryness using a Vacufuge centrifugal concentrator. Dried peptides were stored at -80°C until further use.

### Enrichment and Fractionation of Crosslinks by Size Exclusion Chromatography

Crosslinked peptides derived from *E. coli* extracts and in-cell crosslinked samples were enriched by peptide size exclusion chromatography (SEC). Dried peptides were reconstituted in 250 µL of 3% acetonitrile, 0.1% TFA with vigorous sonication. Peptides were fractionated on an AKTA Go FPLC (Cytiva) with a Superdex 200 10/300 GL gel filtration column. Chromatography used an isocratic method with Solvent A (3% acetonitrile, 0.1% TFA) and a flow rate of 0.4 mL/min (approximate column pressure 2.5 MPa) with a column temperature of 4 °C. Prior to fractionation, the column was equilibrated with Solvent A for 2 hours. 250 µL peptides were injected onto the column after being loaded into a 500 µL loop, and 0.2 column volumes (the void volume) were passed to waste without being collected. Elution volumes were set to 0.5 mL for fractions 1–12, and 1.0 mL for fractions 12–24. Out of 24 fractions, 4–20 were found to be enriched with crosslinked peptides, and these fractions were selected for LC-MS/MS analysis. A_280_ traces from the SEC enrichment or proteome-wide crosslinking experiments can be found in Figure S9. Fractions were reduced to dryness in a Vacufuge centrifugal concentrator and stored at -80°C until further use.

### LC-MS/MS

This study utilized an UltiMate3000 UHPLC system (Thermo Fisher) coupled with a Q-Exactive HF-X Orbitrap mass spectrometer (Thermo Fisher). Peptide fractions were resuspended in 30 µL of 0.1% Optima formic acid (FA) in Optima water, vigorously vortexed, and sonicated for five minutes. The amount of peptide material in each resuspended sample was approximately quantified with a NanoDrop One^C^ microvolume UV-Vis spectrophotometer (Thermo Fisher Scientific), and typically ∼1 µg of the peptide was injected onto the LC-MS system for each run. If a fraction contained insufficient material to inject one µg of peptide, 10 µL of the sample (one-third) was injected instead. The column and trap cartridge temperatures were set to 40 °C, and the flow rate was set to 300 µL/minute. Mobile phase Solvent A comprised 0.1% FA, 2% Optima acetonitrile (Fisher Scientific), 98% Optima water. Mobile phase Solvent B comprised of 0.1% FA in 98% Optima acetonitrile, 2% Optima water (Fisher Scientific).

The injected sample was loaded onto a trap column cartridge (Acclaim PepMap 100, C18, 75 µm × 2 cm, 3 µm, 100-Å column) and washed with Solvent A for 10 minutes. Next, the trap column was switched to be in line with the separating column (Acclaim Pepmap RSLC, C18, 75 µm × 25 cm, 2 µm, 100-Å column). The sample was separated in a linear gradient from 2% B to 35% B over 100 minutes, to 40% B over 25 minutes, and to 90% B over 5 minutes. The end of the LC method employed a saw-tooth gradient to remove sample residue from the resolving column.

XL-MS data were acquired in positive ion mode with each full MS scan using a mass range of 350–1500 *m/z*. The resolution of the full MS scans was set to 120k, the automatic gain control (AGC) target was set to 3E6, the maximum injection time (IT) was set to 100 ms, and a stepped HCD fragmentation scheme using a three-tiered stepped normalized collisional energy (NCE) scheme. Various NCEs were tested to optimize these parameters for the different experiment types and documented in Tables S1-4; however, 22%, 25%, 28% proved optimal for most of the experiments (Supplementary Information). After each full MS scan, ten data-dependent MS^2^ scans were collected at a resolution of 15k, an AGC target of 2E5, a minimum AGC target of 8E3, a maximum IT of 250 ms, an isolation window of 1.0 *m/z* (some lysozyme samples used an isolation window of 1.4 and 2.0 *m/z*; see Table 1), and a dynamic exclusion time of 60.0 s. Ions with charges less than +3 or greater than +8 and isotopomeric peaks were excluded from data-dependent MS^2^ scans.

### XL-MS Data Analysis Workflow

Raw files were converted to the mzML format as centroid spectra using the MSConvert GUI from the ProteoWizard toolkit,^45^ and a spectral search for crosslinked peptides was conducted with MeroX^46^ (version 2.0.1.4). For both DizSEC and DizSPC, protease cleavage sites were allowed after Arg and Lys, with Lys cleavage blocked by Pro, and a maximum of three missed cleavages were allowed. The minimum peptide length was set to 3 residues, and the maximum peptide length was set to 30 residues. Static modifications included the carbamidomethylation of cysteine (C to B), and variable modifications included a maximum of 1 oxidation of methionine (M to m) per peptide (greater than 1 prohibitively expands the search space). The precursor precision was set to 10.0 ppm, and the fragment ion precision was set to 20.0 ppm. Mass recalibration for each sample was determined manually by performing a separate search on each raw file for linear peptides in Proteome Discoverer 2.4 (Thermo Fisher Scientific) without spectrum recalibration, and the median Δ*m/z* (typically between 5–8 Th) was relayed to the MeroX settings file for each individual search. The lower mass limit was set to 350 *m/z*, the upper mass limit was set to 1500 *m/z*, the signal-to-noise ratio was set to 2.0, *b-* and *y-* ions only were enabled, the minimum number of fragments per peptide was set to 3, the minimum MS1 charge was set to 3, and deisotoping mode was selected. We used a pre-score intensity of 10% and an FDR cutoff of 1%. Decoys were generated by shuffling sequences but keeping protease sites within the supplied FASTA files. Typical scores of identified peptides ranged between 20 and 120. Lysozyme samples used quadratic mode, whilst *E. coli* extracts and in-cell crosslinking samples utilized proteome-wide mode developed by Götze and co-workers.^16^ For the lysozyme sample, a FASTA file was prepared that contained the sequence of Hen egg white lysozyme (P00698). For the *E. coli* lysates and in-cell crosslinked samples, the reference FASTA for *E. coli* K-12 substrain MG1655 (UP000000625) was used.

To encode the crosslinkers in MeroX, for the DizSEC-crosslinked samples, the composition mass was set to C_5_NOH_7_ (97.05276383 Da). The first specificity site for the NHS-carbamate was Lys and the N-terminus, and the second specificity site for the diazirine was set to all amino acids, both termini, carbamidomethylated cysteine, and oxidized methionine. Modifications at site 1 were CO with a composition mass of COH_–2_ (25.97926456 Da) and the unmodified peptide. Modifications at site 2 were amine modification C_4_NH_9_ (71.07349927 Da), isocyanate modification C_5_NOH_7_ (97.05276383 Da), and unmodified peptide.

For the DizSPC-crosslinked samples, the composition mass was set to C_6_NOH_9_ (111.06841389 Da). Specificity sites and modifications at site 1 were the same as in the DizSEC settings. Modifications at site 2 were amine modification C_5_NH_11_ (85.08914933 Da), isocyanate modification C_6_NOH_9_ (111.06841389 Da), and unmodified peptide.

### Network Analysis

The analyzed data were exported from MeroX in the XiView format, including identification and sequence files. These files were uploaded to XiView^47^ with a Met oxidation modification mass of 15.99491 Da and a Cys carbamidomethylation modification mass of 57.02146 Da. Each set of experimental data were aggregated together on the XiView server to form network diagrams. Protein Data Bank (PDB) codes were imported into XiView and Euclidian crosslink distance measurements were exported into PyMol scripts. Structures with mapped crosslinks were generated in PyMol by importing the PDB file and running the appropriate crosslink script.

## RESULTS AND DISCUSSION

### Optimization of Crosslinking Parameters on Lysozyme

Initially, we screened several different experimental conditions with DizSEC and DizSPC on a single protein (lysozyme). Parameters screened included concentration of the crosslinker, power of the UV-LED array, presence or absence of a Lys-C pre-digestion step, the addition of DMAP as an acyl-transfer catalyst, and variable collisional energies. The results of these experiments are compiled in Tables S1 and S2, and the crosslinks identified are mapped in Figure S10. Preliminary results confirmed the success of the crosslinking experiment, and crosslinked peptide identification was not very sensitive to the power of the UV lamp, the presence or absence of Lys-C or DMAP, different collisional energies, or even crosslinker concentration— suggesting these compounds function as robust protein crosslinkers. As expected, the identification of crosslinked peptides was explicitly dependent on treatment with a chemical crosslinker and the UV irradiation step (Tables S1-S2). Figure 1D shows an example of an assigned spectrum from MeroX of one crosslinked peptide from lysozyme. The spectrum has 50 peaks that match predicted fragment ions, of which six fragment ions contain the crosslink. Importantly, we detected three of the four fragment ions corresponding to the cleaved crosslink but with no fragmentation of the attached peptides (shown schematically in Figure 1C and annotated in Figure 1D), thereby confirming the generation of an MS-cleavable urea group upon nucleophilic attack by Lys on the NHS-carbamate. Crosslinking results obtained from the DizSEC experimental trials were similar to that of the commercially available non-cleavable photo-crosslinker, sulfo-SDA. Hence, for studies on a single protein, DizSEC could function as an alternative (Table S1). On the other hand, DizSPC had notably fewer crosslinks per trial injection than DizSEC or sulfo-SDA (Table S2).

### A Method for Proteome-Wide Photo-Crosslinking of *E. coli* Extracts

To perform XL-MS on whole extracts from *E.* coli, we used a method illustrated schematically in Figure 2A. Cells were lysed, and lysates were treated with DizSEC/DizSPC, photolyzed, and digested. The complexity of such samples is vast, necessitating a degree of fractionation and enrichment to prevent pervasive co-isolation during LC-MS/MS. We adopted the strategy of peptide size-exclusion chromatography^48^ (alternatives include affinity pull-down on crosslinkers that include an affinity label^49^ or strong cation exchange chromatography^50, 51^). To optimize parameters for identifying crosslinks from proteome-wide photo-crosslinking experiments, we screened several conditions (presence or absence of DMAP, inclusion or exclusion of Lys-C, and HCD energies) and carried out LC-MS/MS on a trial fraction (typically 13 or 14, which tends to be rich in crosslinks; see Tables S3-S4, Figure S9). Based on these data, we decided to move forward with 1 mM DMAP, stepped normalized collisional energies of 22, 25, 28%, and an isolation window of 1.0 *m/z*. In the production set of experiments, these settings were typically applied to fractions 4–19, ca. 16 fractions were separately injected and searched, and the resulting identified crosslinks were pooled together. We performed this experiment four times (a total of 63 LC-MS/MS runs) to generate four distinct datasets, corresponding to crosslinking with either DizSEC or DizSPC and with or without pre-digestion with Lys-C (Figure 2B-C).

As shown in Figure 2B, these crosslinkers generated a satisfactory number of assignable spectra (crosslink-spectrum matches, XSMs). Despite the fact aliphatic diazirine-based crosslinkers are sometimes considered too reactive and primarily quench with water (resulting in quenched, dead-end products), we found in all four cases that dead-end products (Figure 2B, gray) were identified less frequently than spectra in which crosslinks formed between amino acids (black and red). In all cases, the majority were intra-peptidal (red, as many as 9152 in the DizSEC/–Lys-C condition), as expected for crosslinkers with relatively short spacer arms.

**Figure 2.**
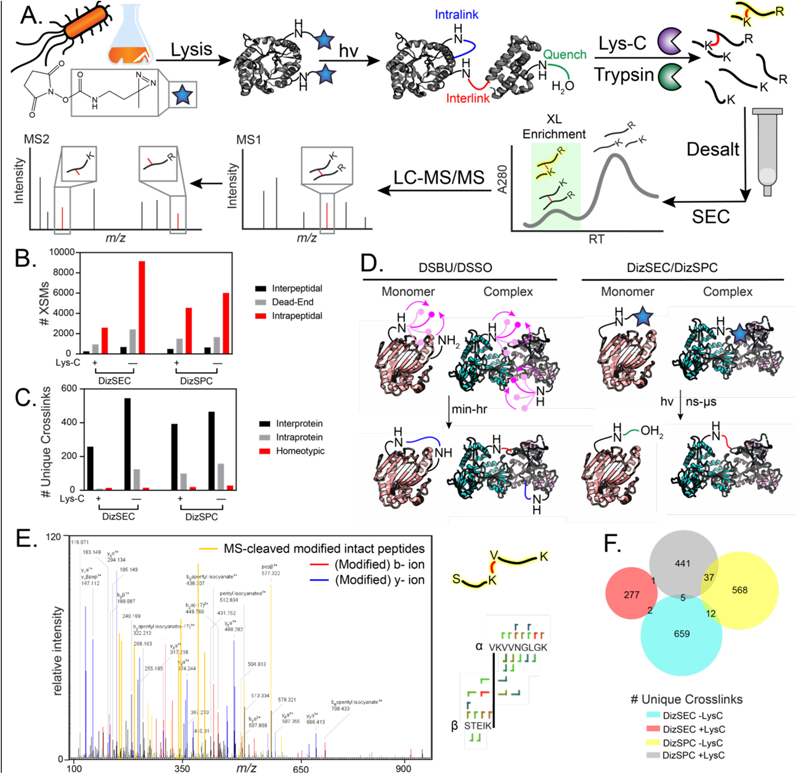
Proteome-Wide Identification of Photo-crosslinks. (**A)** The experimental workflow to prepare crosslinks for LC-MS/MS identification using DizSEC (or DizSPC) on *E. coli* extracts. (**B**) Bar chart showing crosslink-spectrum match (XSM) type distribution. (**C**) Bar chart showing unique crosslink type distribution. (**D**) A model to explain DizSEC/DizSPC’s selectivity for interprotein crosslink formation due to their size and lifetime compared to DSBU/DSSO chemical crosslinking. (**E**) A representative MS2 spectrum from the DizSPC/+Lys-C experiment that matches a crosslinked peptide in which the terminal lysine of one peptide is the attachment site for DizSPC – implying proteolysis at the modified lysine. (**F**) Euler diagram illustrating the general absence of overlap amongst identified crosslinks across the four experimental conditions tested. Figure generated with BioVenn.^52^

Next, we focused on the inter-peptidal XSMs, grouped together redundant ones, and asked among these unique crosslinks, how many bridged across different proteins (interprotein) or within the same protein (intraprotein) (Figure 2C). Strikingly, the majority were interprotein in all conditions (between 75-97%). These findings suggest that DizSEC and DizSPC could be particularly useful for exploring PPIs. However, we were surprised because typically, one sees more *intra*protein crosslinks in most XL-MS studies.^53^ These observations led us to speculate why photo-crosslinkers might be better suited for identifying protein interactions. Figure 2D illustrates our model. With homo-bifunctional chemical crosslinkers, the relatively long persistence time of the NHS-ester moiety enables the crosslinker to “wag” around its protein after initial attachment, exploring the surface, until it eventually finds a suitable partner (another nucleophilic residue). On the other hand, in a hetero-bifunctional photo-crosslinker, the activated carbene has a very short lifetime. Thus, the scenario in which the carbene would have to explore around the surface to find a second attachment site is more likely to result in quenching. On the other hand, at a protein-protein interface, the carbene will be buried and more likely to react with the partner protein. Additionally, DizSEC and DizSPC have short spacer arms, and after the initial reaction, they result in relatively small modifications (cf. Figure 1B), akin to post-translational modification on Lys. This modification is less likely to sterically block a PPI compared to most homo-bifunctional chemical crosslinkers, which will still be bulky after initial attachment. Consequently, one can imagine a surface lysine reacting with DizSEC/DizSPC to form a modified lysine, which later gets buried at a PPI and crosslinked during UV irradiation.

Upon searching for crosslinked spectra, we encountered a sizable number in which the crosslinked lysine occupied a terminal position (as high as 1239 in DizSPC/–Lys-C), implying that both trypsin and Lys-C can cleave proteins at carbamylated lysines which form upon acyl transfer with the NHS-carbamate group. Figure 2E shows an example of a fragmentation spectrum of a terminal lysine-linked peptide, highlighting the substantial number (18) of fragment ions containing the crosslink attached to the assigned lysine in question. We infer that many other XL-MS studies could increase the number of identified crosslinks if the search criteria were adjusted to allow for such linkages. For all four experimental conditions, 24.1% of unique crosslinks were bonded to acidic residues, consistent with a previous report which found acidic residues to be common attachment sites for diazirines.^28^

Finally, we observed that the unique crosslinks across these conditions were mostly unique (Figure 2F), implying that different crosslinkers or proteolysis conditions can be combined to increase coverage. It is also likely a result of the stochasticity of data-dependent acquisition of such complex samples.

### Protein Interaction Networks Derived from Photo-crosslinks

Pooling together the data obtained on all four experimental crosslinking conditions on *E. coli* extracts, we identified 1986 unique crosslinks (1636 interprotein and 350 intraprotein), 1882 unique crosslinked proteins, and 1617 PPIs, some of which include putative or uncharacterized proteins. Figure 3A illustrates a high-level view of this protein interaction network. Details are compiled in Tables S5–S8.

**Figure 3.**
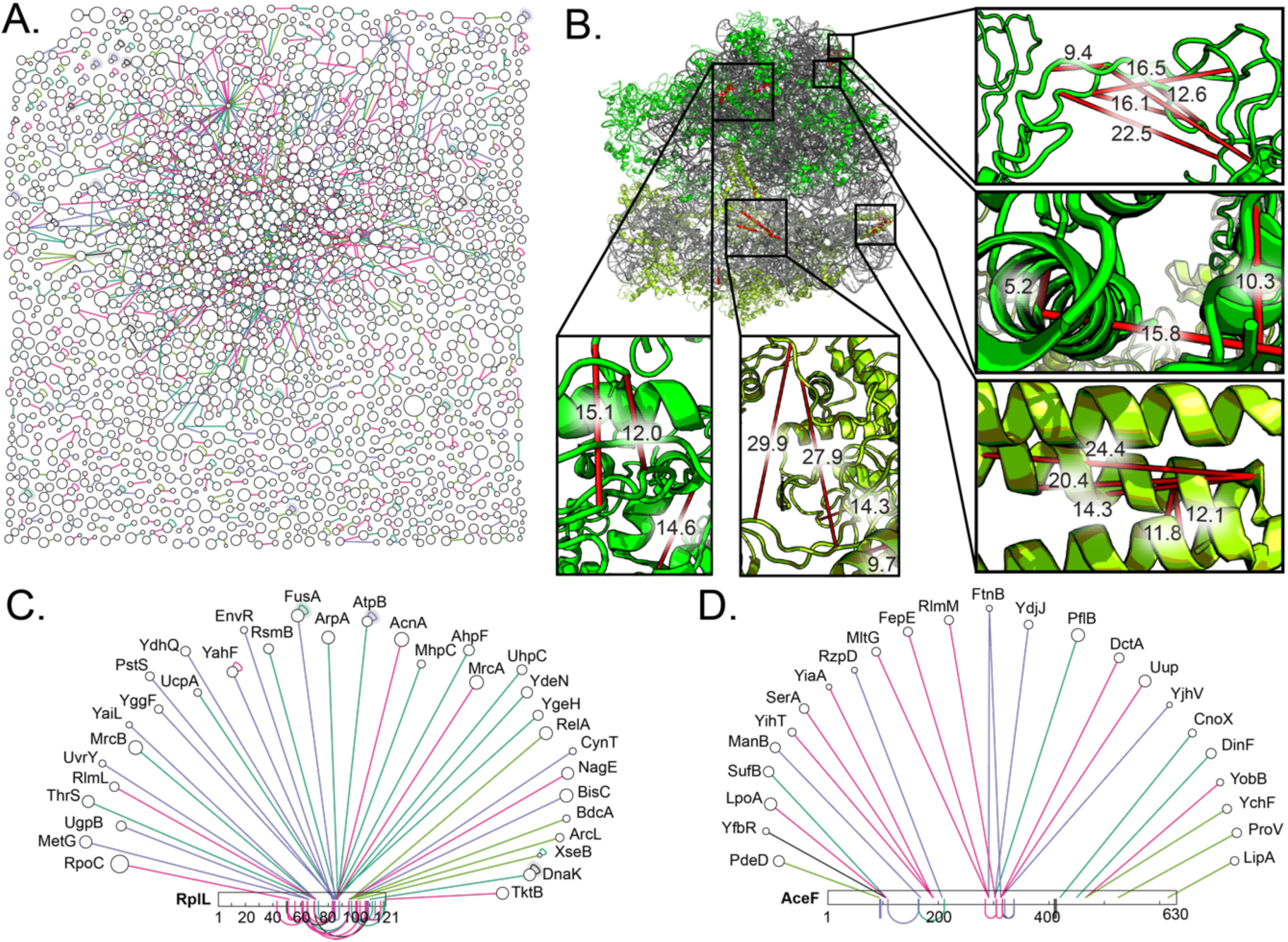
A Residue-Level Protein-Protein Interaction Network of *E. coli* Lysate. (**A**) A network plot representing 1986 distinct crosslinks by pooling data from all four experimental conditions on *E. coli* extracts. The inquisitive reader can interrogate this network interactively at bit.ly/DizSEC_1 through the web-based XiView tool.^47^ (**B**) Identified crosslinks associated with ribosomal proteins are in excellent agreement with a cryo-EM structure of the 70S *E. coli* ribosome (PDB: 4V9D).^54^ Five zoom-in views highlight that all Cα–Cα distances are less than 30 Å in Euclidean distance. (**C, D**) Interaction networks of two *E. coli* proteins (RplL (**C**) and AceF (**D**)) that are hubs for interactions with many other *E. coli* proteins. As XL-MS provides information on both protein identity and residue, these networks illustrate regions of the hub proteins that are “hot spots” (65-75, 80-90, and 100-105 on RplL; and 90-105, 190-210, 290-315 on AceF.

The ribosome is among the most well structurally characterized complexes and hence can serve as a fiducial for the reliability of the crosslinks identified (Figure 3B). In total, we observed 21 crosslinks amongst ribosomal proteins, of which 100% had Cα–Cα distances below 30 Å. Figure 3B shows a few regions within the ribosome with a dense set of crosslinks, in which all of the crosslinked residues were within close proximity.

In addition to being a stable complex, the ribosome is a hub for numerous interacting partners, which is also captured in our dataset and highlights a unique advantage of XL-MS over other structural methods. For instance, we found 34 distinct interactors with RplL (including its well-known interaction with the GTPase elongation factor EF-G (FusA),^55^ Figure 3C). *E. coli* RplL is one of the more elusive ribosomal proteins, absent in most x-ray and cryo-EM structures;^56, 57^ our study suggests this may be because it is a dynamic adaptor, introducing a wide array of factors to the ribosome. This includes some anticipated proteins, such as the stringent response initiator RelA (which recognizes deacylated tRNA in the ribosomal A-site^58^), the co-translational chaperone DnaK (Hsp70), several aminoacyl-tRNA synthetases (ThrS and MetG), and methyltransferases that modify the ribosomal RNA (RlmI and RsmG). On the other hand, RplL’s interactome also consists of several uncharacterized enzymes not previously associated with the ribosome (Figure 3C).

Drawing an example from central metabolism, our network demonstrates pyruvate dehydrogenase (AceF) as a hub (Figure 3D). AceF is an important metabolic enzyme that mediates between glycolysis and the citric acid cycle. We see crosslinking between several important metabolic proteins, including phosphoglycerate dehydrogenase (SerA), involved in serine biosynthesis and reduction of α-ketoglutarate to generate 2-hydroxyglutarate.^59^ Another crosslink was observed with a metabolic enzyme lipoate synthase (LipA), which is well-known to interact directly with E2 subunits and is important for AceF functions.^60^ Other than that, AceF also interacted with several uncharacterized proteins, such as a zinc-binding dehydrogenase, YdjJ; a KpLE2 phage-like protein, YjhV; and a carbon-nitrogen hydrolase family protein, YobB.

In both sub-networks, the central proteins are partnered with several putative or uncharacterized proteins. We propose that photo-crosslinking can give insight into their potential roles by capturing their interaction with more abundant and well-understood partners.

### In-cell Proteome-wide Photo-crosslinking

To see if DizSEC and DizSPC were capable of probing PPIs in their native cellular context, we incubated cells with the compounds after washing away the primary amine-rich LB media and directly performed photolysis on live cells. These experiments explicitly took advantage of the high-power UV LED array used in this study, coupled to a flow cell (Figure 4A, see also Experimental Methods) such that each cell is illuminated for approximately 1 s. With such rapid irradiation, we anticipated the results would capture a snapshot of the cell’s “state” during photolysis and trap transient interactions.

**Figure 4.**
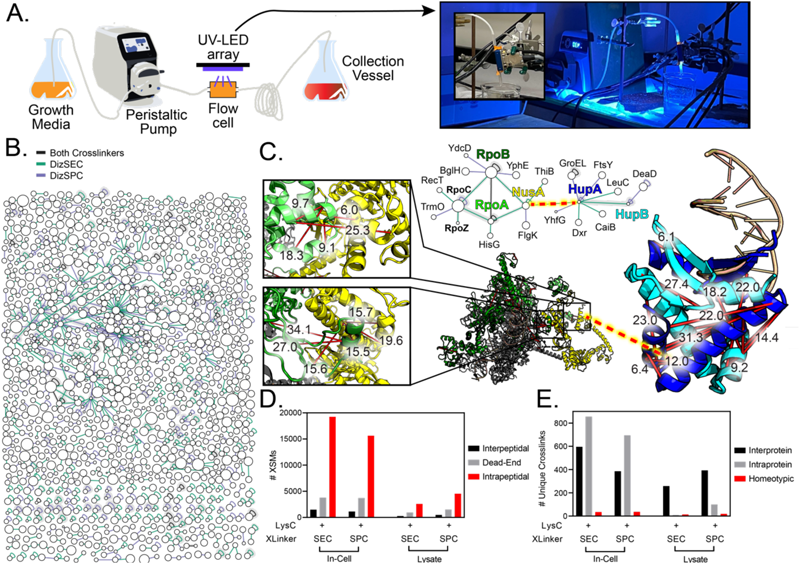
In-cell Photo-crosslinking Visualizes *E. coli* Molecular Sociology at Residue-Level Resolution. (**A**) Workflow for in-cell photolysis. Cells are pumped through a flow cell, where the UV-LED array rapidly crosslinks the samples, which are then collected in a separate vessel. A photo of the setup *in operando* is shown. **(B)** A network plot representing 2131 distinct crosslinks by pooling data from two in-cell photo-crosslinking experiments. The inquisitive reader can interrogate this network interactively at bit.ly/DizSEC_2 through the web-based XiView tool.^47^ **(C)** A subnetwork spanning between the proteins RpoA, RpoB, RpoC, NusA, HupA, and HupB. Crosslinks between RpoA/NusA and RpoB/NusA are mapped onto a cryo-EM structure of the RNA polymerase/NusA complex (PDB: 6FLQ),^61^ and are in excellent agreement. Crosslinks between HupA/HupB are mapped onto an x-ray structure of the HupA/HupB heterodimer (PDB: 4YEW),^62^ and are also in excellent agreement. We additionally discover a crosslink (red dotted-line) between these two structures *in vivo*, putatively involved in transcriptional regulation. **(D)** Comparison of the number and type of XSMs in lysate and in-cell experiments. **(E)** Comparison of the number and type of unique crosslinks identified in lysate and in-cell experiments.

To our delight, in-cell crosslinking studies proved to be an even richer source of identifiable crosslinks. We identified 1486 and 1122 unique crosslinks in-cell *vs.* 281 and 511 unique crosslinks in the lysate (for DizSEC and DizSPC, respectively).

This success might be attributed to the cytosol containing proteins at a higher concentration (>100 mg/mL) and possessing water at a lower concentration than lysates. It also shows that DizSEC and DizSPC are quite cell-permeable, even for a Gram-negative bacterium with two membranes. We point out that whilst SDA is cell-permeable, it is poorly soluble in water, which prompts many users to use sulfo-SDA, an analog that is highly water-soluble but not cell-permeable.^63^ Our results show that such a trade-off does not exist for DizSEC and DizSPC: they are both reasonably soluble in water (when pre-dissolved in DMSO and diluted) and cell-permeable.

### Protein-Protein Interactions *in vivo* Captured in 3-D

We pooled data from two in-cell experimental crosslinking conditions (DizSEC and DizSPC, both with Lys-C pre-digestion) and obtained 14,941 XSMs and 2131 unique crosslinks spanning across 1243 unique proteins. The result is a partial interaction map of the *E. coli* cytosol with residue-level resolution (illustrated schematically in Figure 4B). One example is a sub-network centering on RNA polymerase (Figure 4C), whose core enzyme in *E. coli* is a tetramer, [(RpoA)_2_(RpoB)(RpoC)]. The XL-MS data recapitulate the key interactions that make up the core enzyme, with crosslinks bridging between all three polypeptides. We further identified a linkage between RpoC and RpoZ, the ω-subunit of RNA polymerase, which is an assembly factor that has been shown to facilitate the incorporation of RpoC into the holo-complex.^64^

A previous XL-MS/cryo-ET study in *Mycoplasma* showed that NusA is an adaptor protein that connects RNA polymerase with ribosomes to constitute the “expressosome,” the complex responsible for coupled transcription-translation.^5^ Our results show that the C-terminal helix of NusA binds against a hairpin between the first two alpha helices of HupA, a highly abundant subunit of the HU histone-like DNA binding complex. HU is a heterodimer containing one monomer of HupA and HupB, and is implicated in nucleoid organization and regulation.^65^ Because XL-MS data provide spatial resolution for these linkage sites, this crosslink could be used as distance restraint for future studies focusing on the structural basis of transcriptional regulation *in vivo*. Moreover, these observations cement the role of NusA as an adaptor between RNA polymerase and its various interactors.

To further illustrate the power of our approach with one final example, we mapped out residue-level interaction networks of selected components of the *E. coli* proteostasis network (Figure 5). It is commonly accepted that chaperones have a diverse array of interactors (e.g., DnaK has been shown to have over 650 clients by co-precipitation^66^), promoting the folding of a wide array of clients. Consistent with this view, in our unbiased photo-crosslinking-based dataset, we find that chaperones indeed have a higher number of interactors than average. Because the coverage in our experiments is spread thin over the whole cell’s interactome, these networks are not comprehensive, but they represent a starting point that could be expanded upon by coupling our approach with pull-down enrichment following cell lysis.

**Figure 5.**
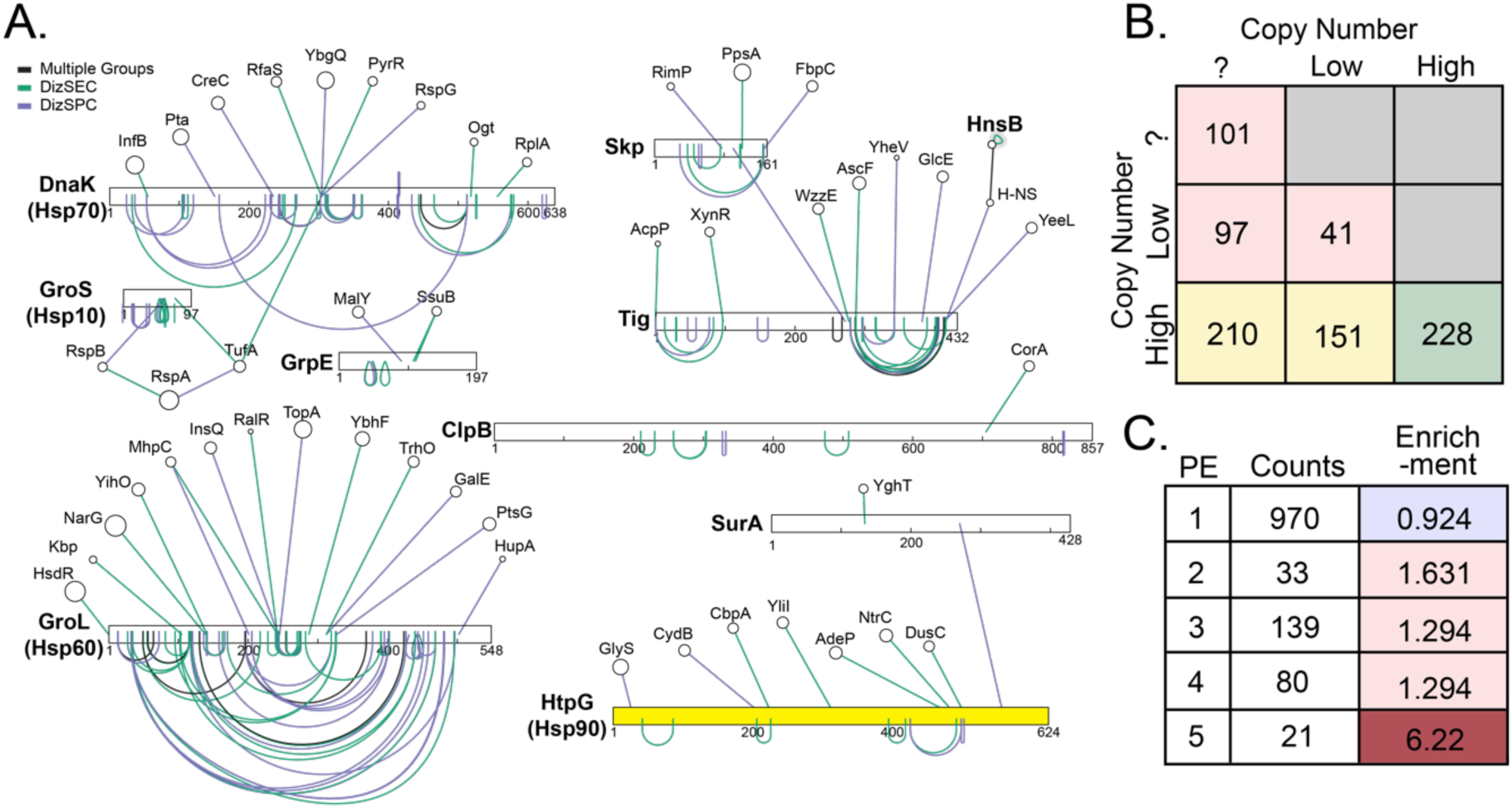
Photo-crosslinks of the *E. coli* proteostasis machinery. (**A**) Chaperone-client crosslink networks derived from the in-cell crosslinking experiments, demonstrating the ability of XL-MS to capture interactions, including those with a cryptic chaperone, HtpG (highlighted in yellow). (**B**) Number of unique inter-protein crosslinks detected in in-cell photo-XL-MS experiments, categorized by the abundance of the two proteins connected by the crosslink. Low-abundance proteins are defined as those with ≤ 200 copies/cell.^73^ Proteins without annotated abundance in ref. 73 are treated as low abundance; green (high-high), yellow (high-low); red (low-low). (**C**) Number of proteins identified in in-cell photo-XL-MS experiments, categorized by their Protein Existence (PE) score. Enrichment factors are calculated relative to search on linear peptides.

In particular, the XL-MS strategy introduces a method to identify clients and their interaction sites with the more elusive *E. coli* chaperone, HtpG (Hsp90). Chaperones that tightly bind their substrates (such as DnaK) or fully encapsulate them (such as GroL/S) have been particularly amenable to pull-down based proteomics to exhaustively catalog their client substrates.^66, 67^ These methods have been more challenging to apply to HtpG, and consequently our knowledge of its substrate scope is significantly more limited.^68, 69^ We note a crosslink between HtpG with the periplasmic chaperone, SurA suggests an intriguing link between the cytosolic and periplasmic proteostasis networks. The crosslink is within the HtpG substrate-binding clamp suggesting that HtpG might be involved in promoting the biogenesis of SurA,^70^ which could explain why HtpG-null knockouts are deficient in their ability to process several periplasmic proteins, β-lactamase and alkaline phosphatase.^71^

### Shedding Light on the “Dark” Proteome

We were surprised to identify many crosslinks between chaperones and proteins that are either low abundance or putative/ uncharacterized (Figure 5A). Since chaperones tend to be highly abundant molecules within the cell, we theorized that crosslink formation between high abundance and low abundance proteins could make it easier to identify peptides from low abundance proteins. We revisited the full in-cell photo-XL-MS dataset, and for each unique interprotein crosslink, assessed whether it bridged between two high-abundance proteins (copy number > 200; roughly the upper two-thirds of the proteome^72, 73^), two low-abundance proteins (copy number ≤ 200; the bottom third of the proteome), or between a high- and low-abundance protein. Indeed, we found an unusually high number of crosslinks involving low-abundance proteins (Figure 5B, yellow). Moreover, we found a high number of crosslinks involving proteins with an unreported copy number, which can be qualitatively interpreted as “low-abundance” given the high coverage of the Ribo-Seq experiments on which the abundance data were based.^72, 73^ The frequency of proteins appearing in this dataset that were not covered by Ribo-Seq is striking, given that proteomics methods typically have lower coverage than those employing Illumina sequencing.

Finally, we sought to characterize how “uncharacterized” the in-cell photo-XL-MS dataset is. For model organisms, UniProt proteome files provides a Protein Existence (PE) score for each protein, in which 1 is the highest (protein is experimentally validated) and 5 is the lowest (protein uncertain). Our network included 1243 proteins in total, of which 21 were in the most uncharacterized category. This represents a 6-fold enrichment (Figure 5C) relative to how frequently proteins in this category are observed in a typical search for linear peptides. Indeed, all the PE categories greater than 1 were enriched in our dataset, a difference that was statistically significant by the chi-square test (P < 10^-24^). We theorize that these uncharacterized proteins may be enriched by crosslinking to high abundance proteins, allowing us to identify transient or unstable interactions that were not identified with affinity purification mass spectrometry (AP-MS) or other interactomic methods.

## CONCLUSION AND FUTURE DIRECTIONS

Over the past decades, diazirines have achieved a privileged position in chemical biology and chemoproteomics because of their versatility as a “universal” crosslinking moiety. Nevertheless, despite the commercial availability of SDA and sulfo-SDA, they have yet to be broadly exploited for XL-MS studies because of high quenching rates (lowering the yield of crosslinked peptides), their high reactivity and diverse crosslinking sites (complicating spectral search and limiting the software capable of identification), and absence of MS-cleavability. In this study, we demonstrate with the two reported compounds (DizSEC and DizSPC) that by combining diazirenes with NHS-carbamates on a single compound, the complexity associated with identifying photo-crosslinks can be greatly reduced by utilizing the MS-cleavability of the urea functional group formed upon acyl transfer to protein targets. We thereby demonstrate the feasibility of proteome-wide identification of photo-crosslinked peptides. Though it was not a part of the original molecular design, the fortuitous water-solubility and membrane permeability of DizSEC and DizSPC enables such experiments to be conducted *in cellulo*, elucidating protein interactions at residue-level resolution in their native context.

In this study, we identified many PPIs that include proteins with unknown functions that have so far not been ascribed interactors or partners. We speculate that photo-XL-MS may be particularly well-suited for catching such interactions because the photochemistry is very rapid, thereby trapping potentially transient interactions, and because crosslinks formed between low-abundance uncharacterized proteins and their higher-abundance interactors could help overcome the sensitively challenge of the former.

Low-abundance proteins that do not have catalytic activities and are not associated with stable complexes are often challenging to characterize. Even in the highly studied model organism *E. coli*, there is experimental evidence for only 69.3% of the proteome at the protein level and 72.8% at the transcriptional level based on the PE score system.^74^ Based on these early photo-XL-MS experiments, we speculate that the method can enrich for low abundance proteins that interact with high abundance proteins, thereby providing a new tool to assign putative functions to uncharacterized proteins. Remarkably, many proteins that we detected in our proteome-wide search for crosslinked peptides were *not* identified in conventional searches for linear peptides on the same data. This observation is notable because it is common in XL-MS studies to minimize the protein database employed for the crosslink search to include only the proteins that were found in a linear search – the wisdom being that this will ease the more complex crosslink search. In contrast, we show that by combining the efficient MS-cleavability of DizSEC/DizSPC-derived crosslinks with the recently developed “proteome-wide mode” in MeroX,^16^ protein database restriction is neither necessary nor desirable, enabling our photo-XL-MS study to identify proteins that were otherwise unidentifiable.

We anticipate photo-XL-MS could have two potential broader impacts on molecular bioscience. As we have emphasized, photo-crosslinks yield information on structure (at the residue-level of resolution) as well as interaction. Recent years have seen increased interest in characterizing the quinary structure of proteins,^75, 76^ which refers to the specific sticking interactions that cause the cytosol to demix and enable particular sets of proteins to have higher local concentrations and adopt preferential orientations with respect to one another. Exascale computing has enabled molecular dynamics simulations to be conducted on systems approaching the size of cells,^77, 78^ and with these “computational microscope” studies the structure of transient protein assemblies in a cellular milieu are coming into focus. To date however, there are few experimental techniques that can measure quinary interactions to test the predictions from computational microscopy. We propose that the partial interaction maps based on photo-XL-MS could help refine these models and create dialogue between experiment and simulation.

Similarly, the emerging field of molecular sociology^41, 42, 79^ aims to provide a census of interactions between biomolecules in their native environment and has been driven primarily by the extraordinary advances in cryo-electron tomography (cryo-ET). These studies have so far focused on large molecular complexes with distinct shapes (such as the ribosome^80^ or proteasome) and in bacteria with very spacious cytosols, such as *Mycoplasma pneumoniae.*^15^ One current roadblock is that whilst the spatial resolution of cryo-ET is remarkable and improving, its “compositional resolution” – that is, the ability to identify which specific protein an arbitrary segment of electron density corresponds to – is more limited. In this regard, XL-MS and cryo-ET have complementary strengths (compositional and structural resolution, respectively) and hence a close collaboration between them is set to enable molecular sociology to break new ground toward characterizing smaller proteins that are less easy to identify based on electron density alone. The advent of photo-XL-MS using the tools and methods we have presented here should be particularly germane toward this effort.

## ASSOCIATED CONTENT

### Supporting Information

All the additional data of crosslinking experiments, synthetic procedures, and characterization of synthesized compounds. Spreadsheets listing the pooled unique crosslinks identified in the lysate and in-cell crosslinking experiments are provided as Data S1 and Data S2.

### Data Deposition

Raw data corresponding to the crosslinking experiments in extracts (63 LC-MS/MS runs in total) and to the in-cell crosslinking experiments (31 LC-MS/MS runs in total) have been deposited to PRIDE under accession code XXXX. Also included are MeroX method files (.mxf) and MeroX output files (.zhrm).

## AUTHOR INFORMATION

### Notes

The authors declare no competing financial interest.

## ACKNOWLEDGEMENTS

S.D.F. acknowledges support from the NIH Director’s New Innovator Award (DP2GM140926). P.S. would like to acknowledge support from the Albstein Foundation for

Brain Research. The authors would like to thank Yingzi Xia for bringing the novel NHS-carbamate chemistry from the Chamot-Rooke Lab to our attention and for developing the SEC fractionation method for crosslinking enrichment. The authors also thank Dr. Phil Mortimer for providing critical mass spectral support.

## SUPPORTING INFORMATION

### Section 1: Synthetic Methods

All the reagents and chemicals for the synthesis were procured from commercial suppliers and used without further purification. Rf values were determined using TLC silica gel G60 aluminum sheets and visualized by UV at wavelength 254 nm or KMnO_4_. NMR spectra were recorded using Bruker Avance 400 MHz FT-NMR spectrophotometer and chemical shifts are reported as δ (ppm) values.

### Scheme S1: Synthesis of DizSEC and DizSPC^a^

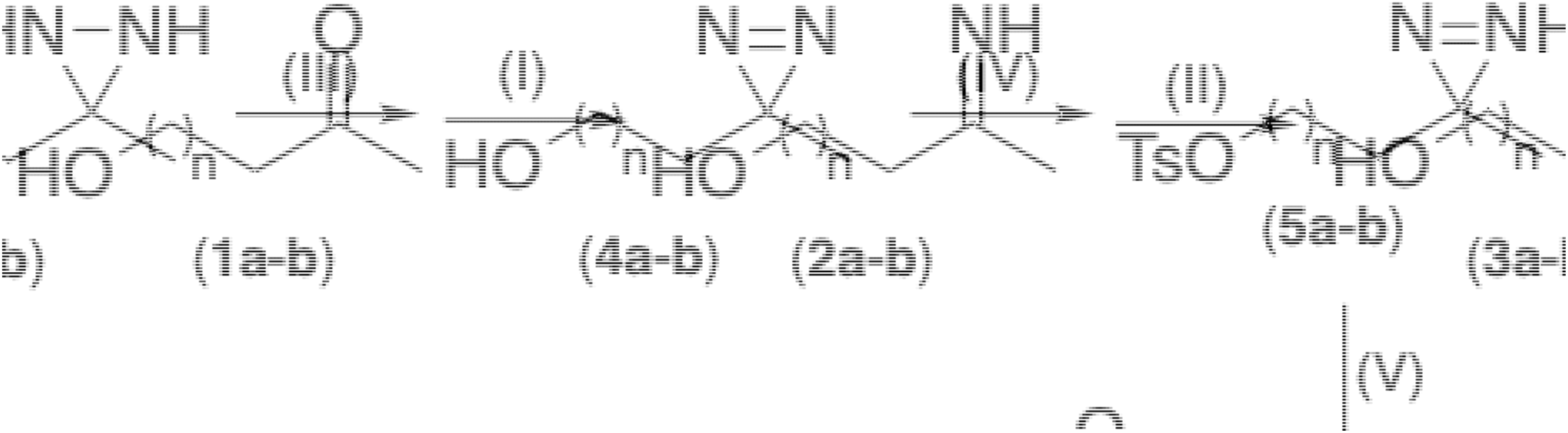

^a^Reagents and conditions: (I) liq. NH_3_ in MeOH, -20 °C, stirring, 3 h; (II) NH_2_OSO_3_H, MeOH, - 20 °C to RT, stirring, overnight; (III) I_2_, Et_3_N, MeOH, 0 °C to RT, stirring, 2 h; (IV) p-TsCl, pyridine, 0 °C to RT, stirring, 2 h; (V) NaN_3_, DMF, RT, stirring, overnight; (VI) PPh_3_, THF:H_2_O (9:1); RT, 6 h; (VII) disuccinimidyl carbonate, CH_3_CN, 0 °C to RT, stirring, 2 h.

#### 2-(3-methyl-3*H*-diazirin-3-yl)ethyl 4-methylbenzenesulfonate (5a) and 3-(3-methyl-3*H*-diazirin-3-yl)propyl 4-methylbenzenesulfonate (5b)^1,2^

In an argon-filled and flame-dried 3-neck round bottom flask, 4-hydroxy-2-butanone (**1a**, 3.0 g, 34.5 mmol, 1 eq)) or 5-hydroxy-2-pentanone (**1b**, 3.5 g, 34.5 mmol, 1 eq) was cooled to -20 °C, and 7N methanolic ammonia solution (35 mL) was added very slowly. The reaction mixture was stirred at -20 °C for 3 h under argon to get respective imines (**2a** or **2b)**.

The solution of hydroxylamine-O-sulfonic acid (4.3 g, 37.9 mmol, 1.1 eq) in anhydrous methanol (25 mL) was added dropwise with a syringe pump to the above reaction mixture containing compound **2a** or **2b** at -20 °C. The reaction mixture was slowly allowed to warm to room temperature and stirred overnight. The insoluble white precipitate was filtered away and washed successively with anhydrous methanol. The filtrate was concentrated *in vacuo* to get respective diaziridine derivatives (**3a** and **3b**).

Compound **3a** or **3b** was re-dissolved in anhydrous methanol (25 mL). The solution was cooled to 0 °C, followed by the addition of triethylamine (14.4 mL, 103.5 mmol, 3 eq). Solid iodine was then added in small portions until the dark brown color of iodine persisted for 10 min. The reaction mixture was warmed to room temperature and stirred for 2 h. The solution of the reaction mixture was concentrated to dryness *in vacuo*, and the concentrated residue was taken in ethyl acetate (50 mL). The organic layer was successively washed with 1N HCl, 10% sodium thiosulfate, and brine. The organic mixture was then dried over anhydrous sodium sulfate and concentrated *in vacuo* to get crude diazirine derivatives (**4a** or **4b**) as a pale yellow oil and used for the next step without further purification.

The resultant compound **4a** (1.3 g, 13.141 mmol) or **4b** (1.5 g, 13.1 mmol) was dissolved in anhydrous pyridine (10 mL), and an equimolar quantity of p-toluenesulfonyl chloride (2.5 g, 13.1 mmol) was added portion-wise at 0 °C. The reaction was allowed to warm at room temperature and stirred for 2 h. The completion of the reaction was monitored by TLC using hexane:ethyl acetate (4:1) as the mobile phase. The reaction mixture was diluted with ethyl acetate and was washed successively with 1N HCl, saturated aqueous sodium bicarbonate, and brine. The organic mixture was then dried over anhydrous sodium sulfate and concentrated *in vacuo*. The crude product was further purified by column chromatography (5-10% ethyl acetate in hexane) using silica gel to get the pure compound as a colorless oil (compound **5a**, 1.7 g, compound **5b**, 1.8 g, 19.7% overall yield for four steps).

#### Compound 5a

^1^H NMR (400 MHz, CDCl_3_) δ_H_ 7.84 (d, *J* = 8.3 Hz, 2H), 7.38 (d, *J* = 8.3 Hz, 2H), 3.97 (t, *J* = 6.4 Hz, 2H), 2.48 (s, 3H), 1.69 (t, *J* = 6.4 Hz, 2H), 1.02 (s, 3H).

#### Compound 5b

^1^H NMR (400 MHz, CDCl_3_) δ_H_ 7.78 (d, *J* = 8.3 Hz, 2H), 7.36 (d, *J* = 8.0 Hz, 2H), 3.99 (t, *J* = 6.1 Hz, 2H), 2.45 (s, 3H), 1.54 – 1.50 (m, 2H), 1.42 – 1.38 (m, 2H), 0.97 (s, 3H).

**Figure S1.**
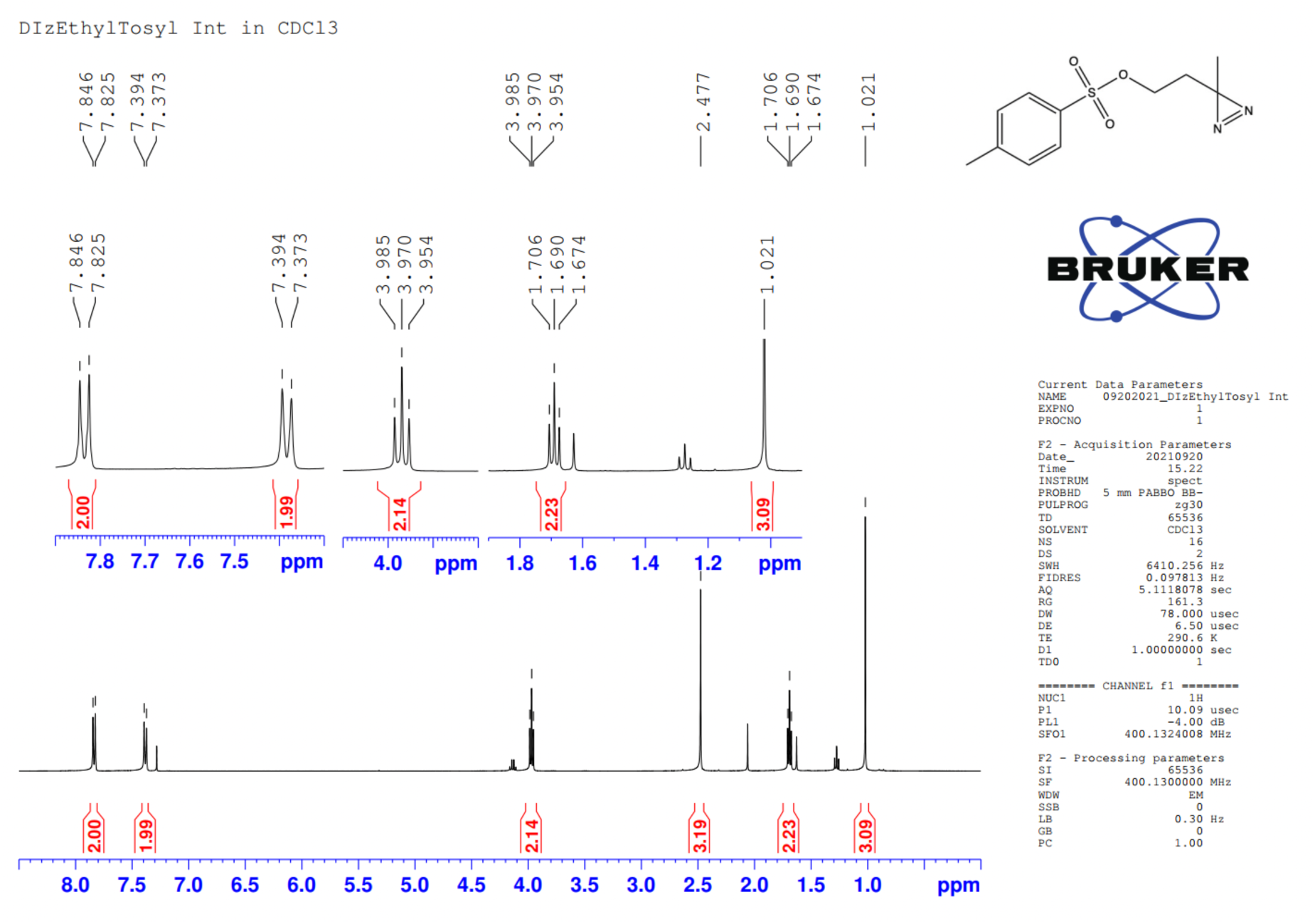
^1^H NMR spectrum of compound **5a**.

**Figure S2.**
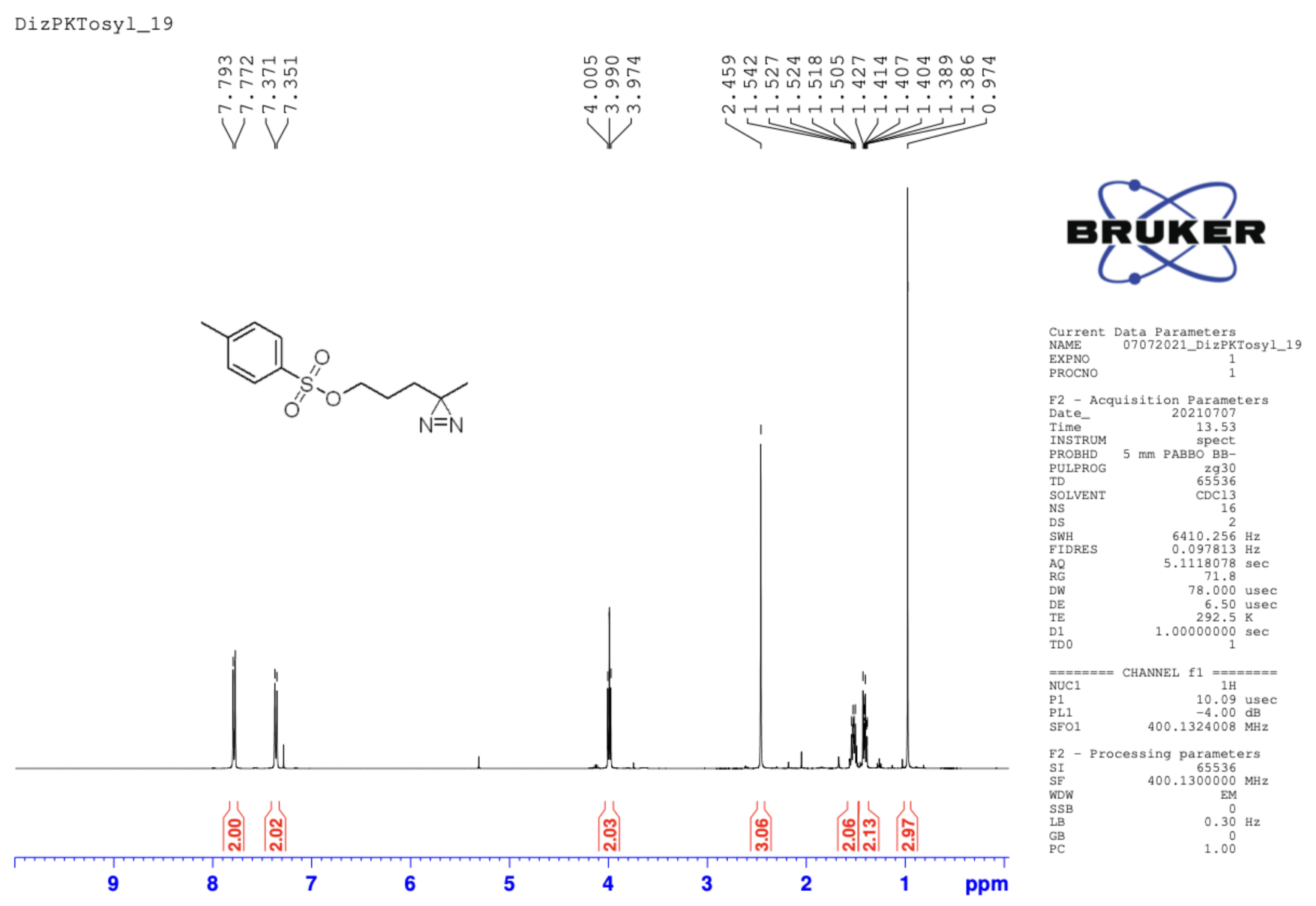
^1^H NMR spectrum of compound **5b**.

#### 2-(3-methyl-3*H*-diazirin-3-yl)ethan-1-amine (7a) and 3-(3-methyl-3*H*-diazirin-3-yl)propan-1-amine (7b)^3^

To the mixture of compound **5a** (1.7 g, 6.8 mmol, 1 eq) or **5b** (1.8 g, 6.8 mmol, 1 eq) in DMF (20 mL), sodium azide (1.5 g, 23.7 mmol, 3.5 eq) was added, and the reaction mixture was stirred at room temperature overnight under N_2_. The resultant mixture was poured on ice water (20 mL) dropwise and extracted with diethyl ether (2 X 20 mL). The combined organic layer was dried over anhydrous sodium sulfate and concentrated in vacuo to get compound **6a** or **6b**, respectively.

To a solution of compound **6a** or **6b** in THF:water (9:1, 10 mL), triphenylphosphine (3.0 g, 11.5 mmol, 1.7 eq) was added, and the reaction mixture was stirred for 6 h at room temperature. The crude reaction mixture was acidified with 1M HCl under an ice-cold condition and washed with diethyl ether (5 X 20 mL). The resulting aqueous solution was further neutralized with 1M NaOH to pH >10.0 and extracted with diethyl ether (4 X 30 mL). The confirmation of desired amine extraction was performed by TLC using DCM:methanol (4:1) as the mobile phase, and ninhydrin was used for visualization. The combined organic layer was dried over anhydrous sodium sulfate and concentrated *in vacuo* to get the compound as a pale-yellow oil (compound **7a**, 0.494 g, 73.3%; compound **7b**, 0.585 g, 76.2%).

#### Compound 7a

^1^H NMR (400 MHz, CDCl_3_) δ_H_ 2.71 (t, *J* = 6.4 Hz, 2H), 1.46 – 1.42 (m, 2H), 1.09 (s, 3H).

#### Compound 7b

^1^H NMR (400 MHz, CDCl_3_) δ_H_ 2.68 (t, *J* = 6.8 Hz, 2H), 1.43 – 1.38 (m, 2H), 1.36 – 1.31 (m, 2H), 1.03 (s, 3H).

**Figure S3.**
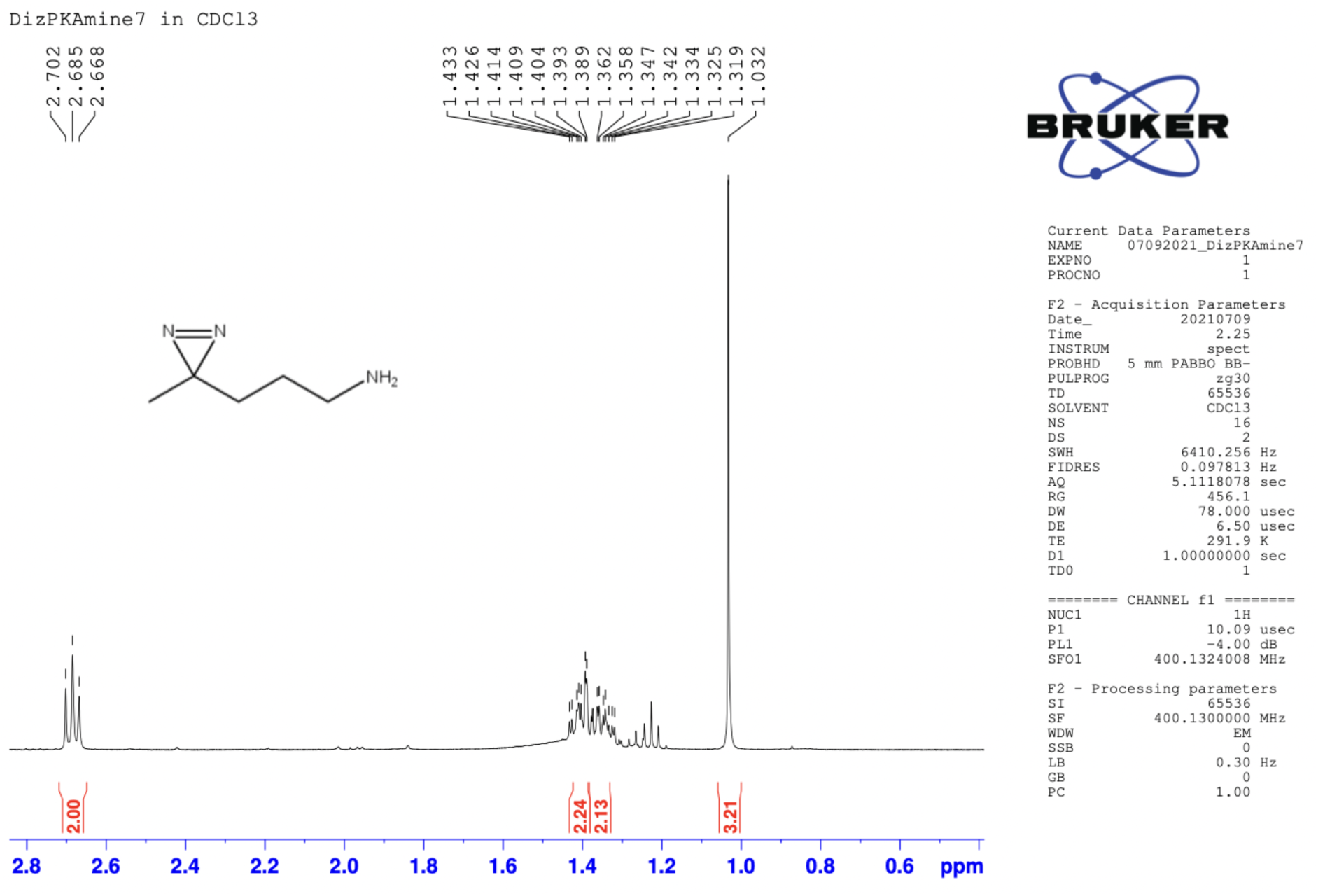
^1^H NMR spectrum of compound **7b**.

#### 2,5-Dioxopyrrolidin-1-yl (2-(3-methyl-3*H*-diazirin-3-yl)ethyl)carbamate (DizSEC, 8a) and 2,5-dioxopyrrolidin-1-yl (3-(3-methyl-3*H*-diazirin-3-yl)propyl)carbamate (DizSPC, 8b)^4–6^

In an ice-cold suspension of disuccinimidyl carbonate (0.204 g, 0.795 mmol, 1.5 eq) in acetonitrile (10 mL), the mixture of compound **7a** (0.053 g, 0.530 mM, 1 eq) or **7b** (0.060 g, 0.530 mM, 1 eq) in acetonitrile (2 mL) was added dropwise with continuous stirring. The reaction was allowed to warm at room temperature and stirred for additional 2 h. The completion of the reaction was monitored by TLC using DCM:MeOH (9:1) as the mobile phase. The reaction mixture was concentrated to dryness *in vacuo*, and the concentrated residue was taken in ethyl acetate (20 mL). The organic layer was successively washed with 10% aqueous citric acid and brine. The organic mixture was dried over anhydrous sodium sulfate and concentrated *in vacuo* to get pure DizSEC (**8a,** 0.111 g, 86.72%) or DizSPC (**8b**, 0.116 g, 85.93%), respectively as white semi-solids.

#### DizSEC (8a)

^1^H NMR (400 MHz, CDCl_3_) δ_H_ 5.45 (s, 1H), 3.20 (dd, *J* = 13.1, 6.9 Hz, 2H), 2.85 (s, 4H), 1.69 (t, *J* = 7.0 Hz, 2H), 1.10 (s, 3H). ^13^C NMR (126 MHz, CDCl_3_) δ_C_ 169.9, 151.4, 53.4, 37.1, 34.0, 25.5, 25.4.

#### DizSPC (8b)

^1^H NMR (400 MHz, CDCl_3_) δ_H_ 5.53 (s, 1H), 3.25 (dd, *J* = 12.6, 6.4 Hz, 2H), 2.84 (s, 4H), 1.56-1.36 (m, 4H), 1.04 (s, 3H). ^13^C NMR (126 MHz, CDCl_3_) δ_C_ 169.9, 151.4, 60.4, 41.4, 31.3, 25.5, 25.3, 19.7.

**Figure S4.**
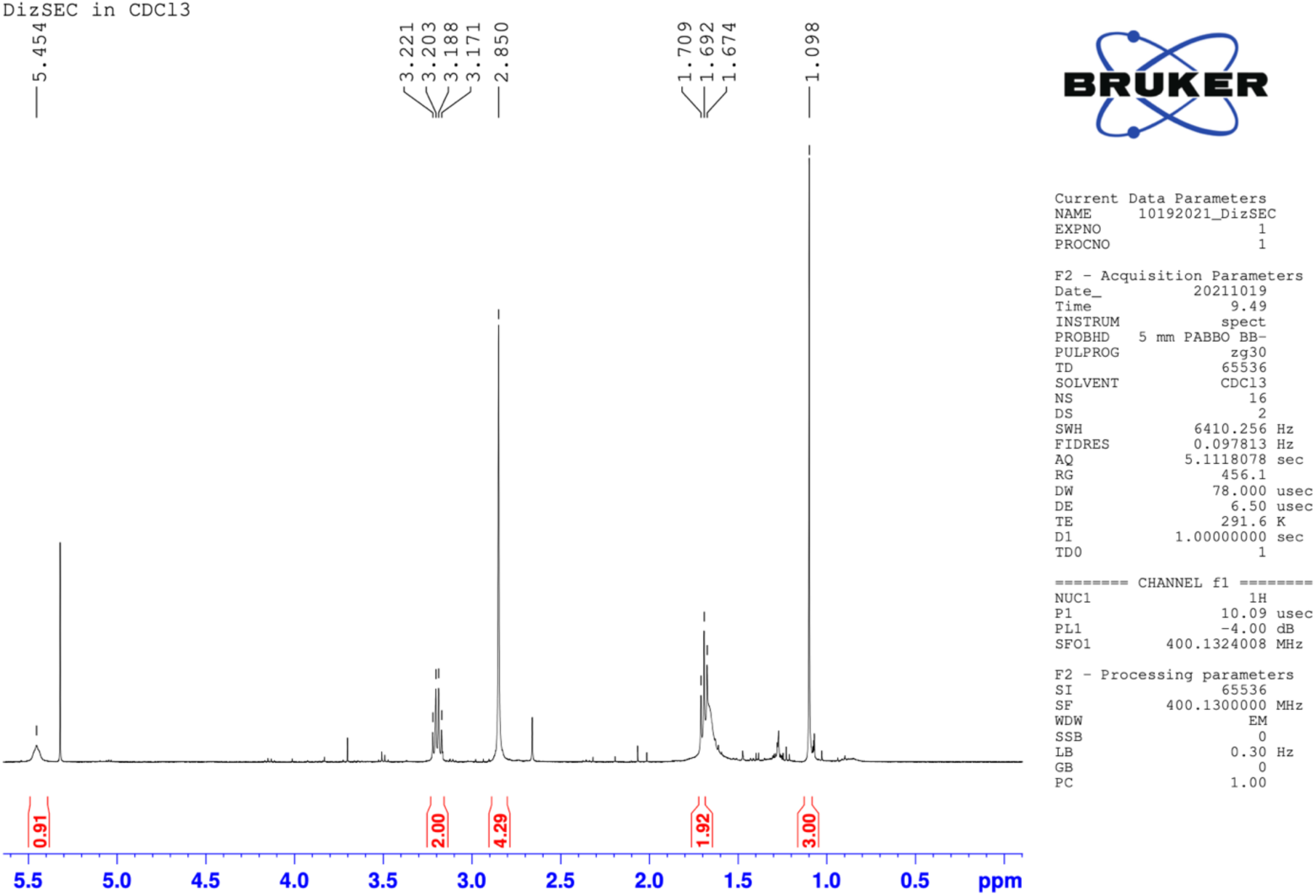
^1^H NMR spectrum of compound **8a**.

**Figure S5.**
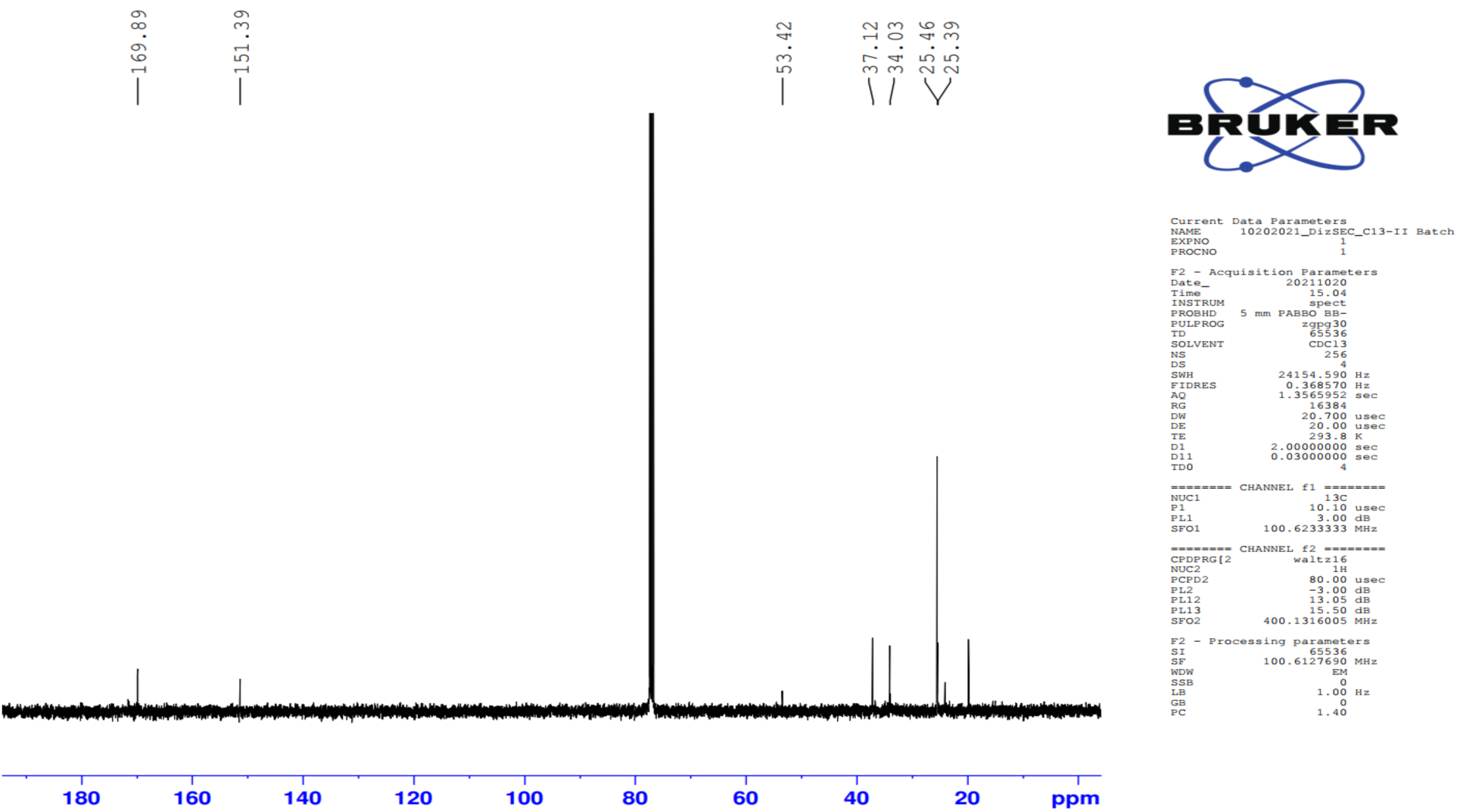
^13^C NMR spectrum of compound **8a**.

**Figure S6.**
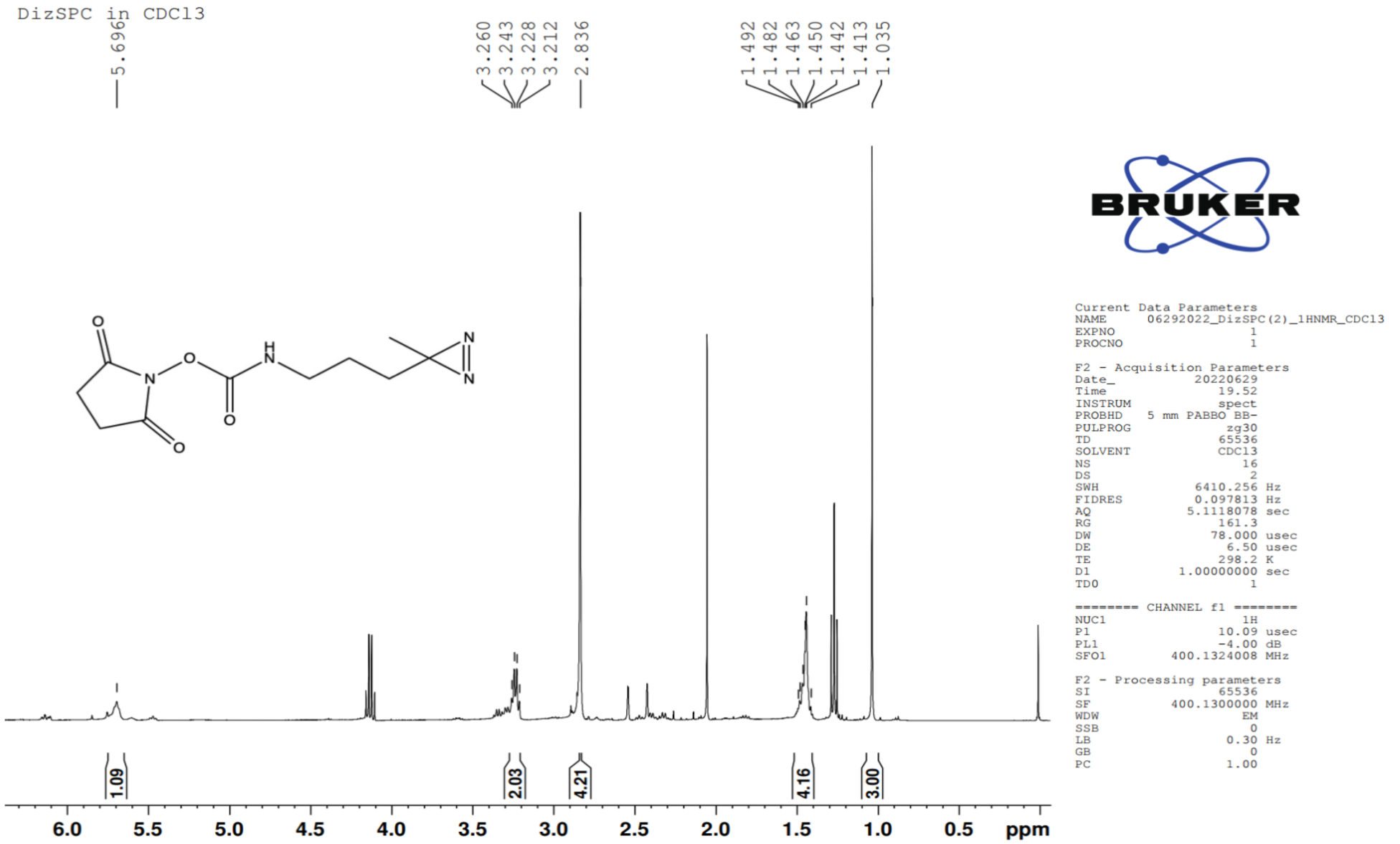
^1^H NMR spectrum of compound **8b**.

**Figure S7.**
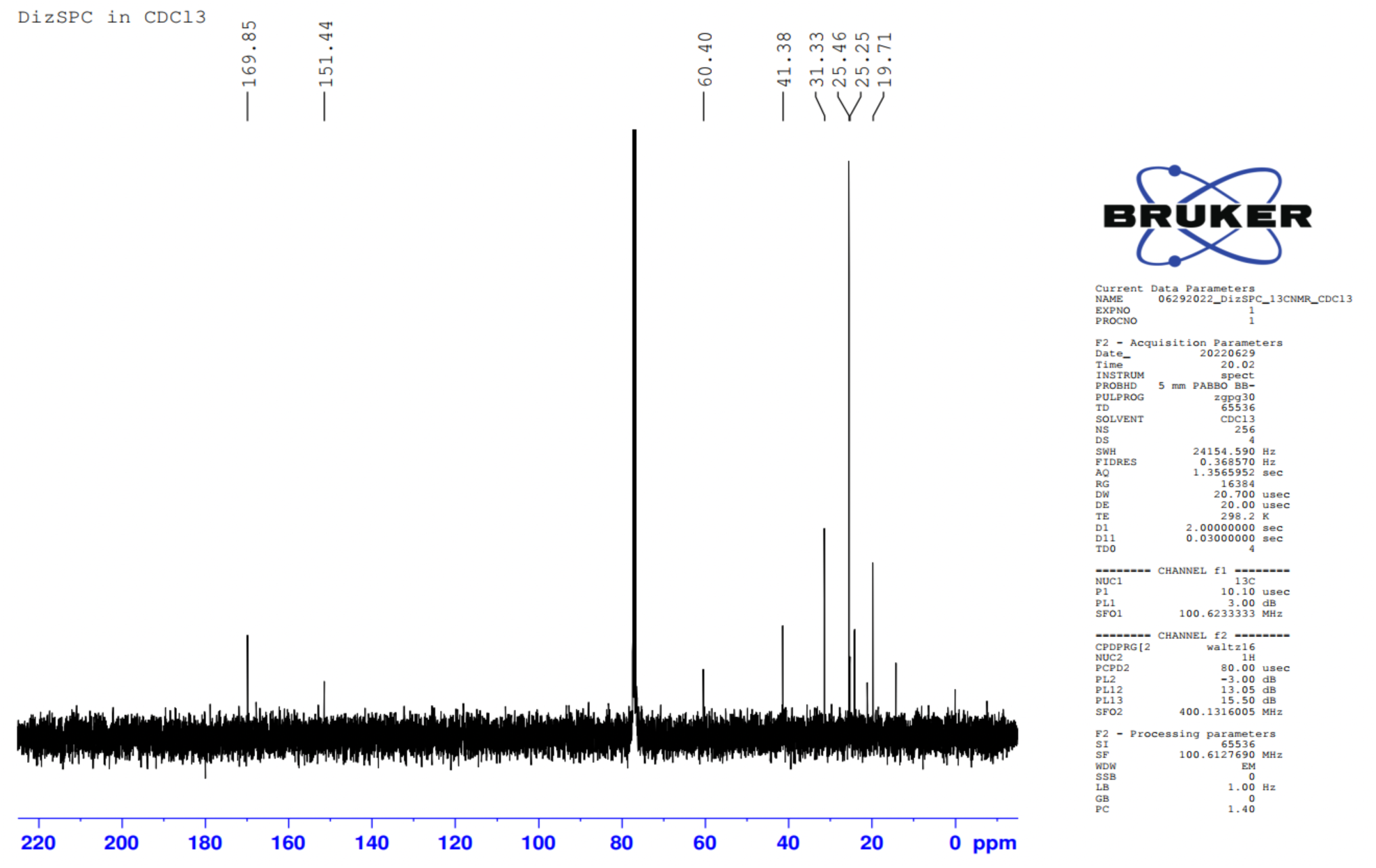
^13^C NMR spectrum of compound **8b**.

### Section 2: Supplementary Figures

**Figure S8.**
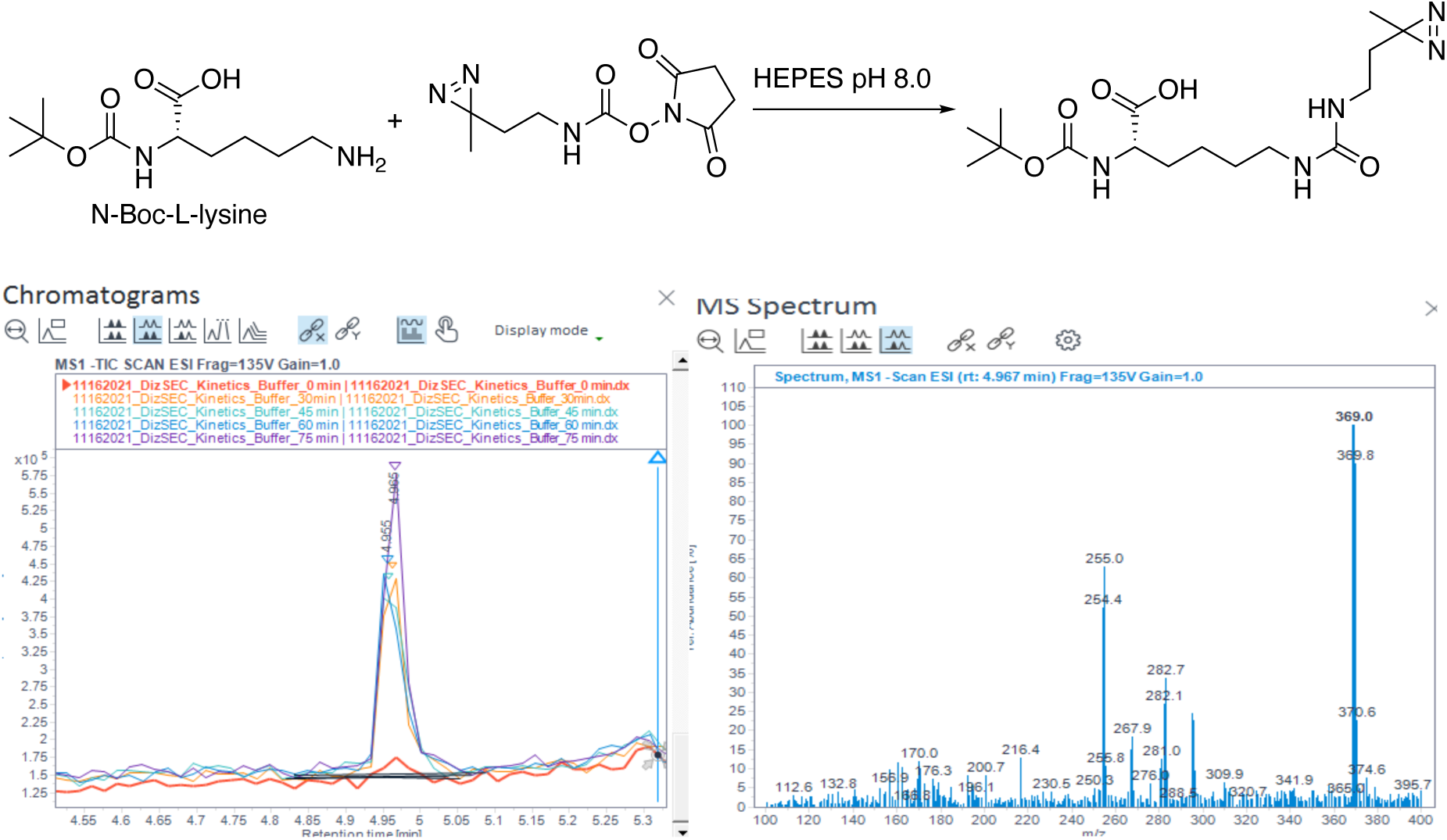
Kinetic Study of DizSEC on Model Compound. *Top*, Reaction used for kinetic study of acyl transfer to NHS-carbamates. *Left,* Overlay of total ion chromatograms of the product, showing the progress of the formation of the urea. *Right,* sample mass spectrum of the product peak. Expected mass of product = 371.

**Figure S9.**
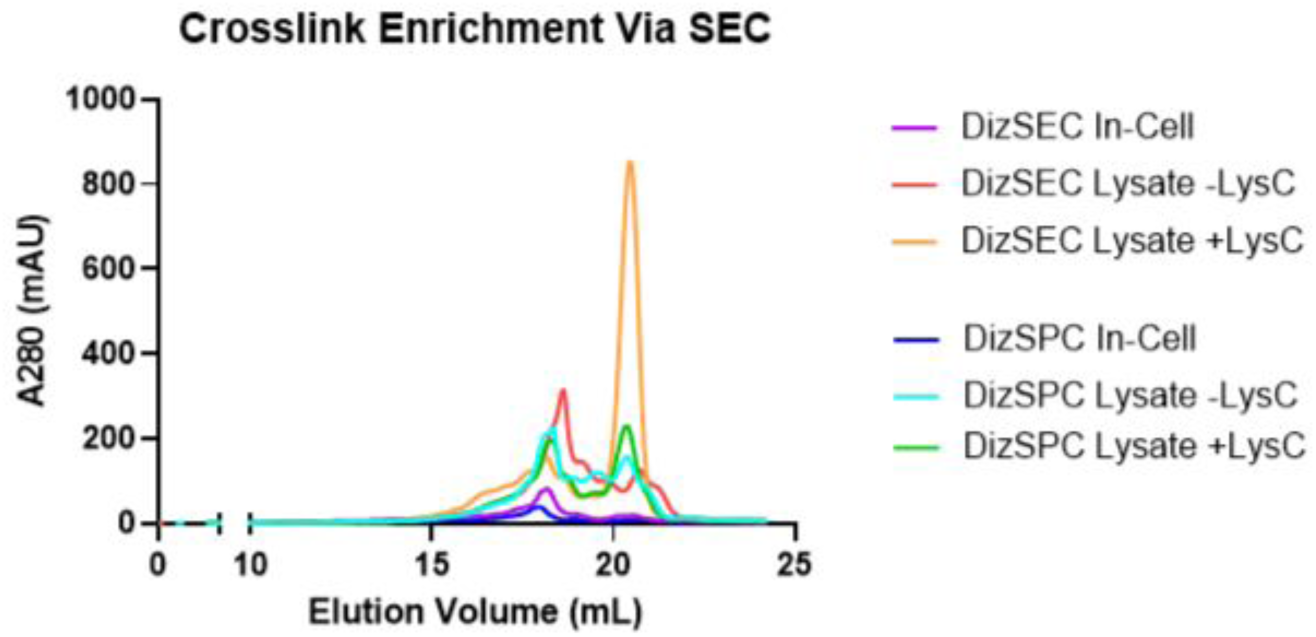
Enrichment of crosslinks using size exclusion chromatography (SEC). Crosslinks were enriched on an Akta Go FPLC system. Crosslinked fractions were collected between 12 mL and 20 mL of elution volume. Between 12 mL and 18 mL of elution volume, up to twelve fractions of 0.5 mL were collected for analysis. Between 18 mL and 25 mL, up to 7 fractions of 1 mL were collected for analysis.

**Figure S10.**
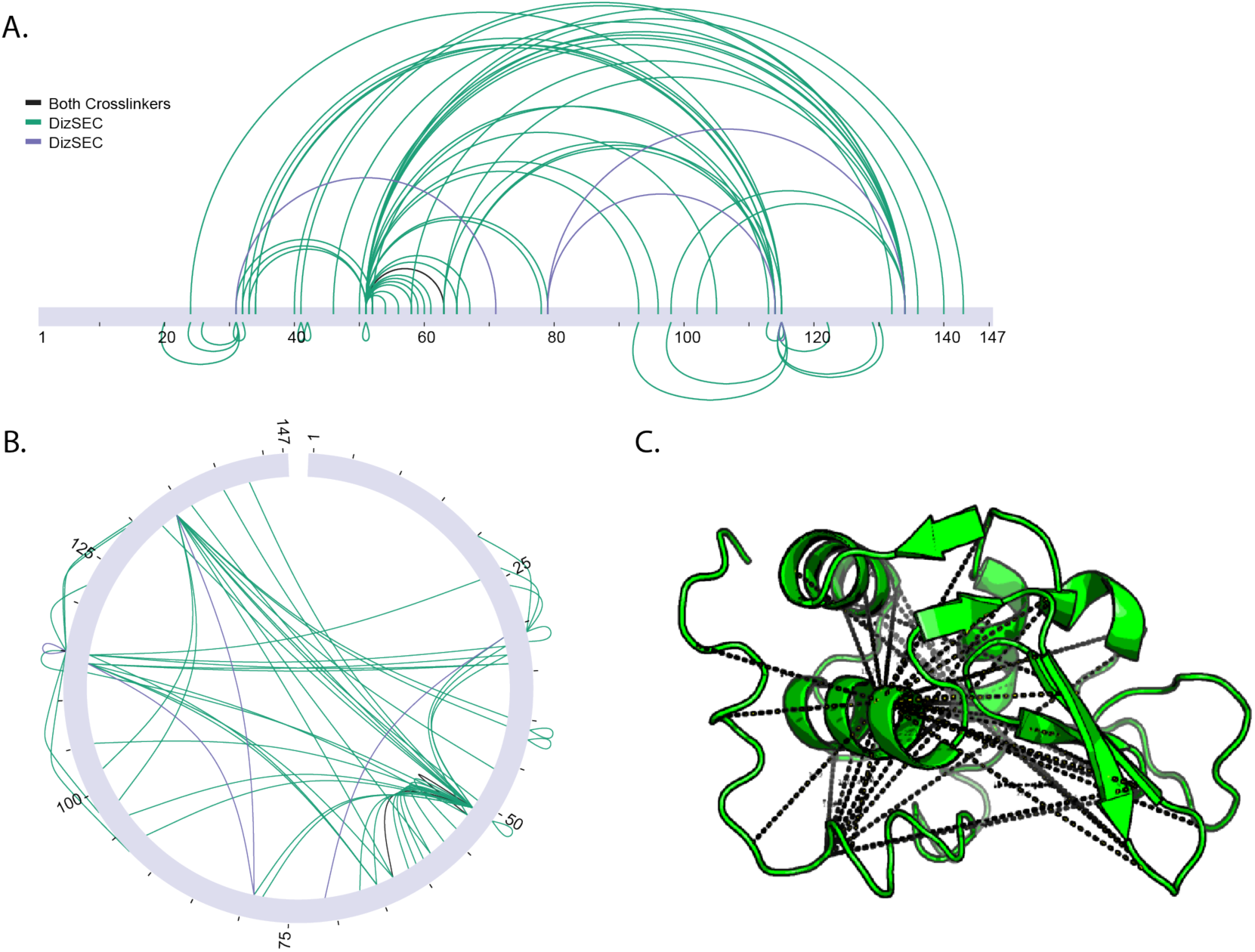
Crosslinking of lysozyme with DizSEC and DizSPC. (A) Crosslinks of lysozyme with DizSEC and DizSPC from all trial experiments shown in linear chart form. (B) Circular chart form. (C) Crosslinks mapped onto a hen-egg lysozyme crystal structure 3LYZ.^7^

### Section 3: Sample MS2 Fragmentation Spectra

**Figure S11.**
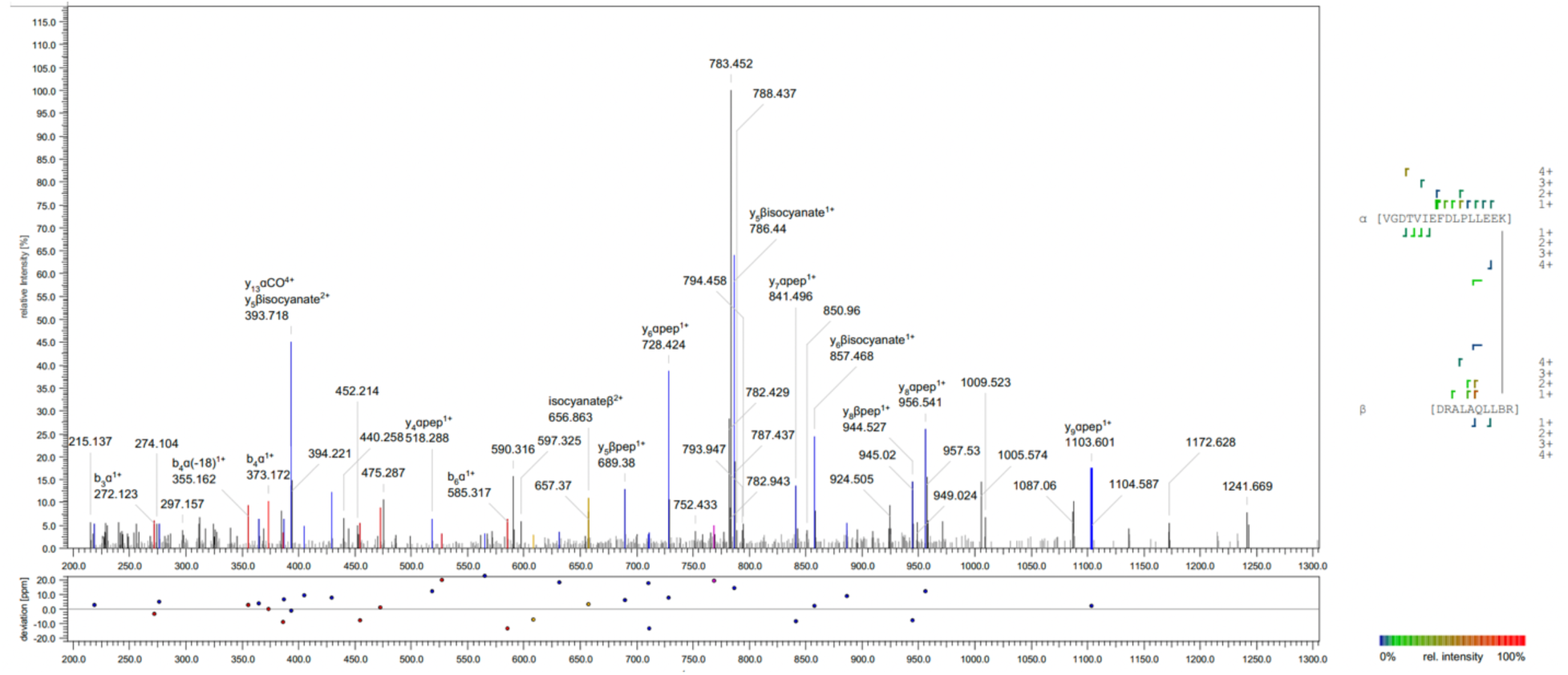
Representative spectrum from the DizSEC lysate experiment predigested with Lys-C. This spectrum shows an interprotein crosslink between the Crr ɑ-peptide [VGDTVIEFDLPLLEEK] crosslinked at position K16 to the SelA β-peptide, [DRALAQLLBR], crosslinked at position B9.

**Figure S12.**
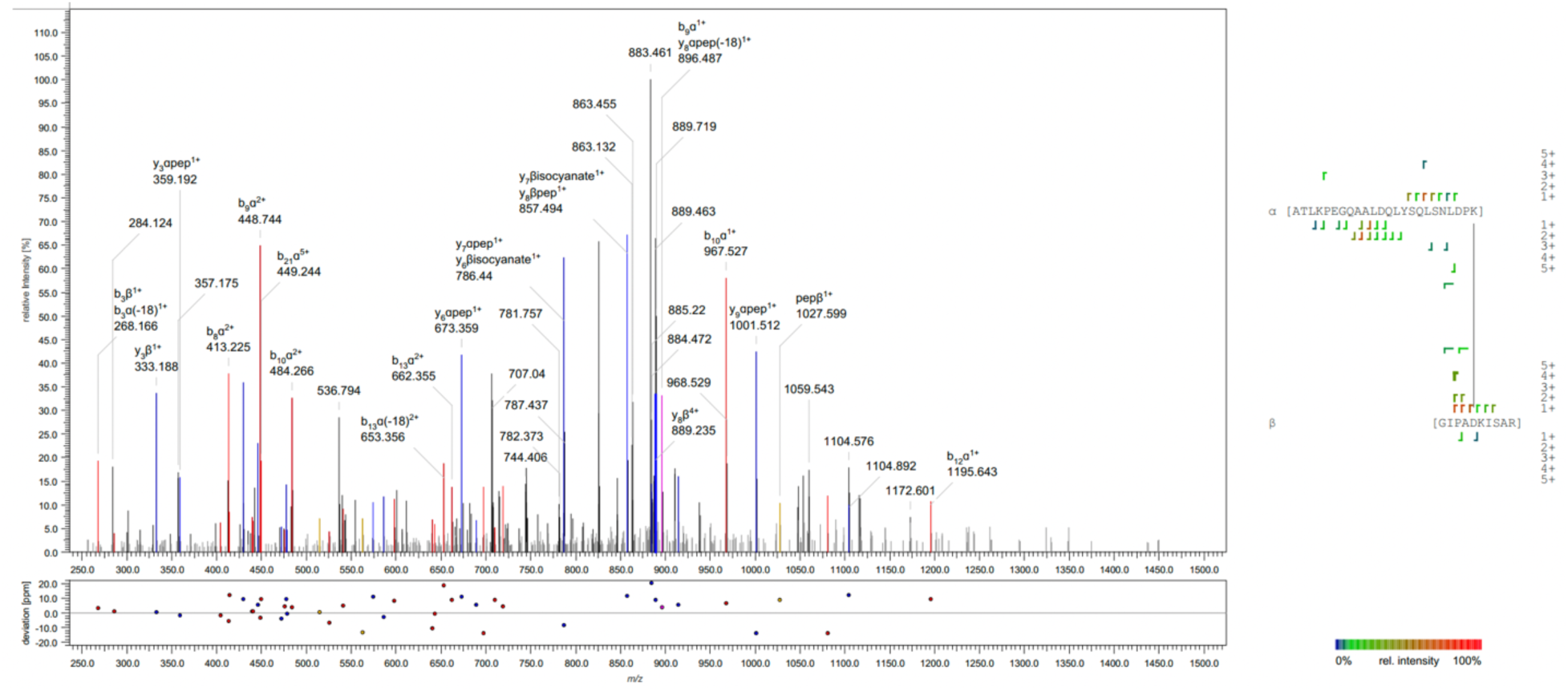
Representative spectrum from the DizSEC lysate experiment predigested with Lys-C. This spectrum shows an intraprotein crosslink in OmpA between the ɑ-peptide [ATLKPEGQAALDQLYSQLSNLDPK] crosslinked at position K24 to the β-peptide, [GIPADKISAR], crosslinked at position D5.

**Figure S13.**
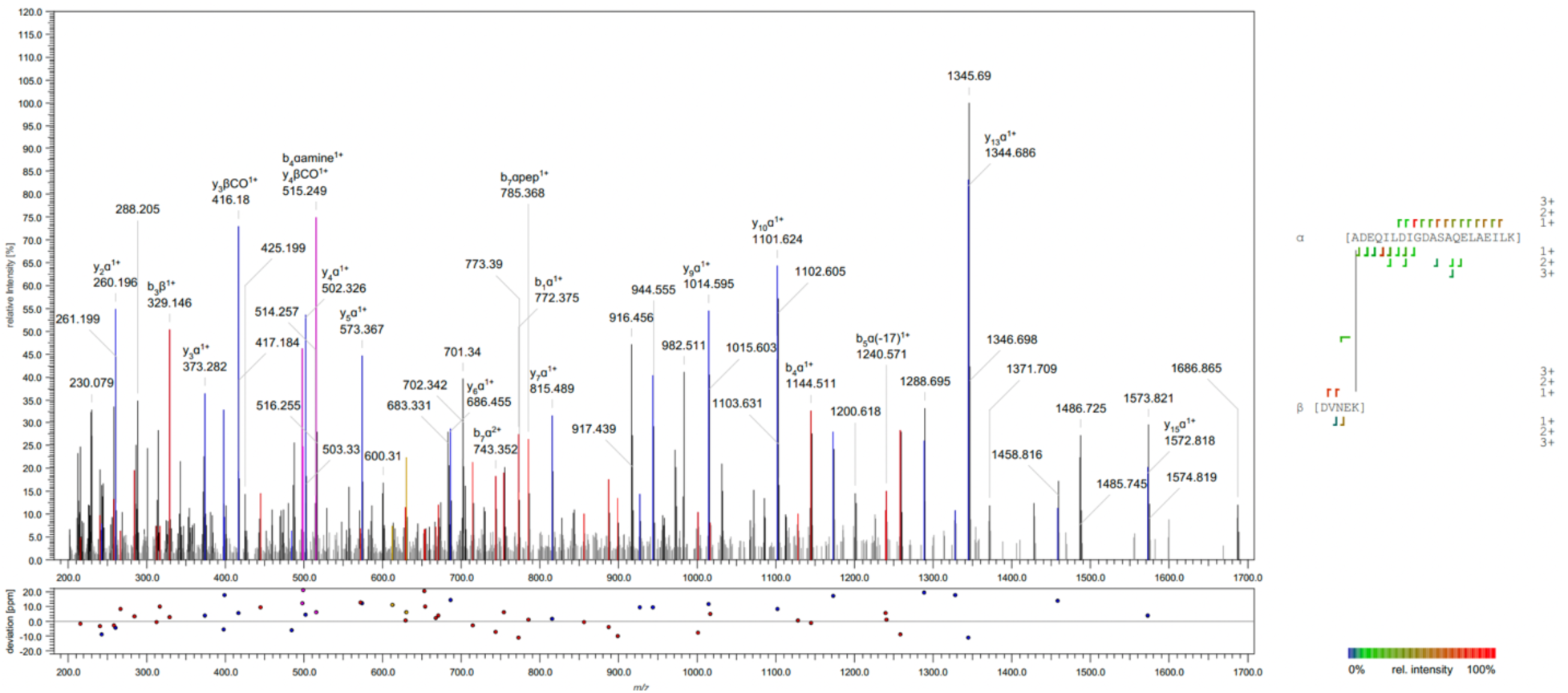
Representative spectrum from the DizSEC lysate experiment without Lys-C. This spectrum shows an interprotein crosslink between the Pgk ɑ-peptide [ADEQILDIGDASAQE-LAEILK] crosslinked at position A1 to the LafU β-peptide, [DVNEK], crosslinked at position K5.

**Figure S14.**
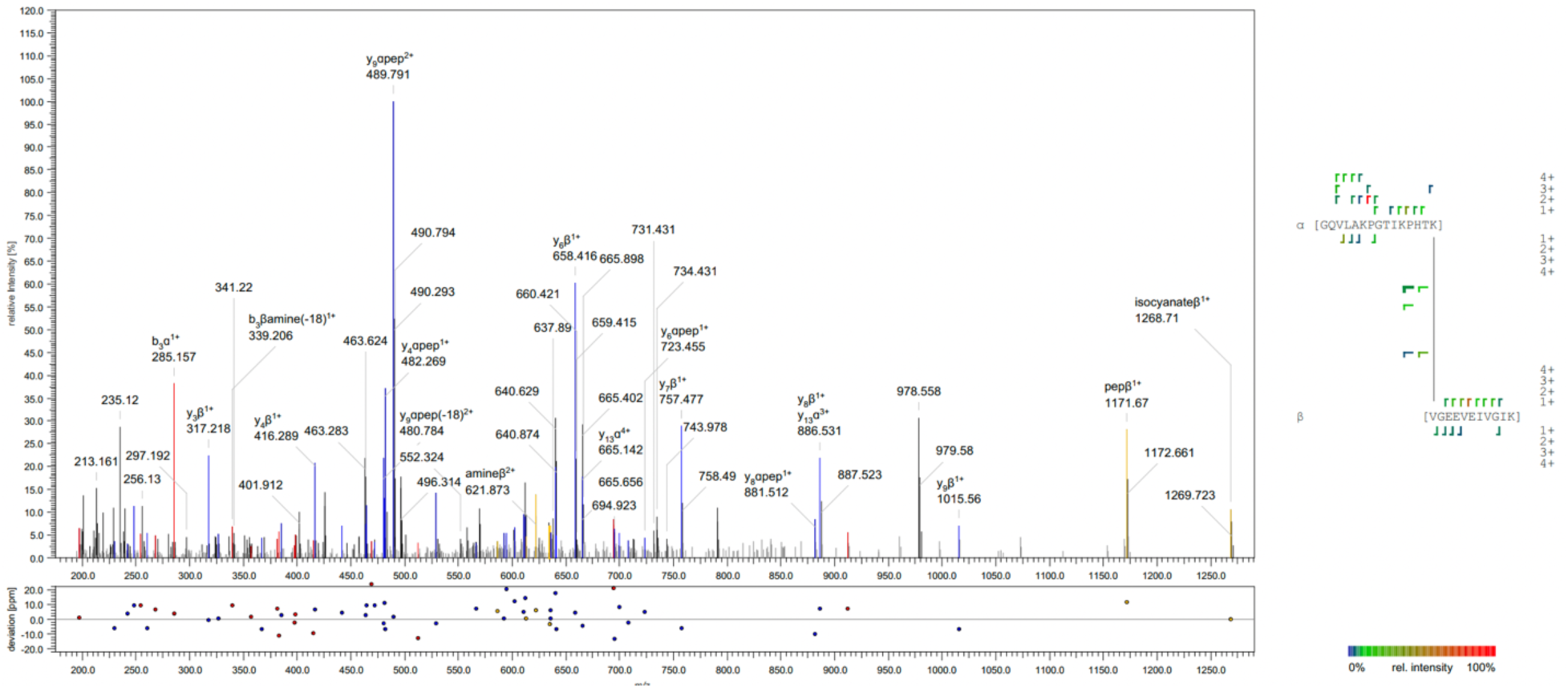
Representative spectrum from the DizSEC lysate experiment without Lys-C. This spectrum shows an intraprotein crosslink in TufA between the ɑ-peptide [GQVLAKPGTIKPHTK] crosslinked at position K15 to the β-peptide, [VGEEVEIVGIK], crosslinked at position V1.

**Figure S15.**
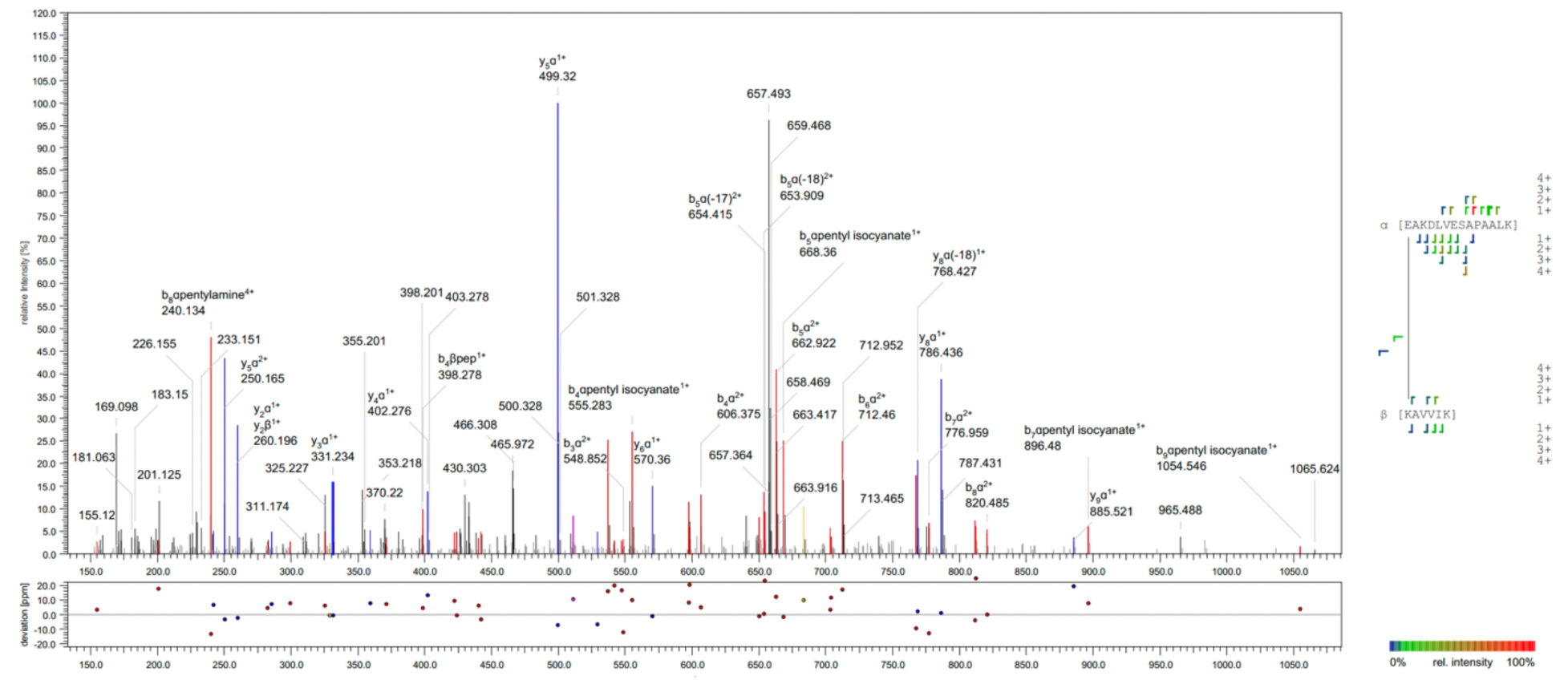
Representative spectrum from the DizSPC lysate experiment predigested with Lys-C. This spectrum shows an interprotein crosslink between the RplL ɑ-peptide [EAKDLVESAPAALK] crosslinked at position E1 to the YgeH β-peptide, [KAVVIK], crosslinked at position K1.

**Figure S16.**
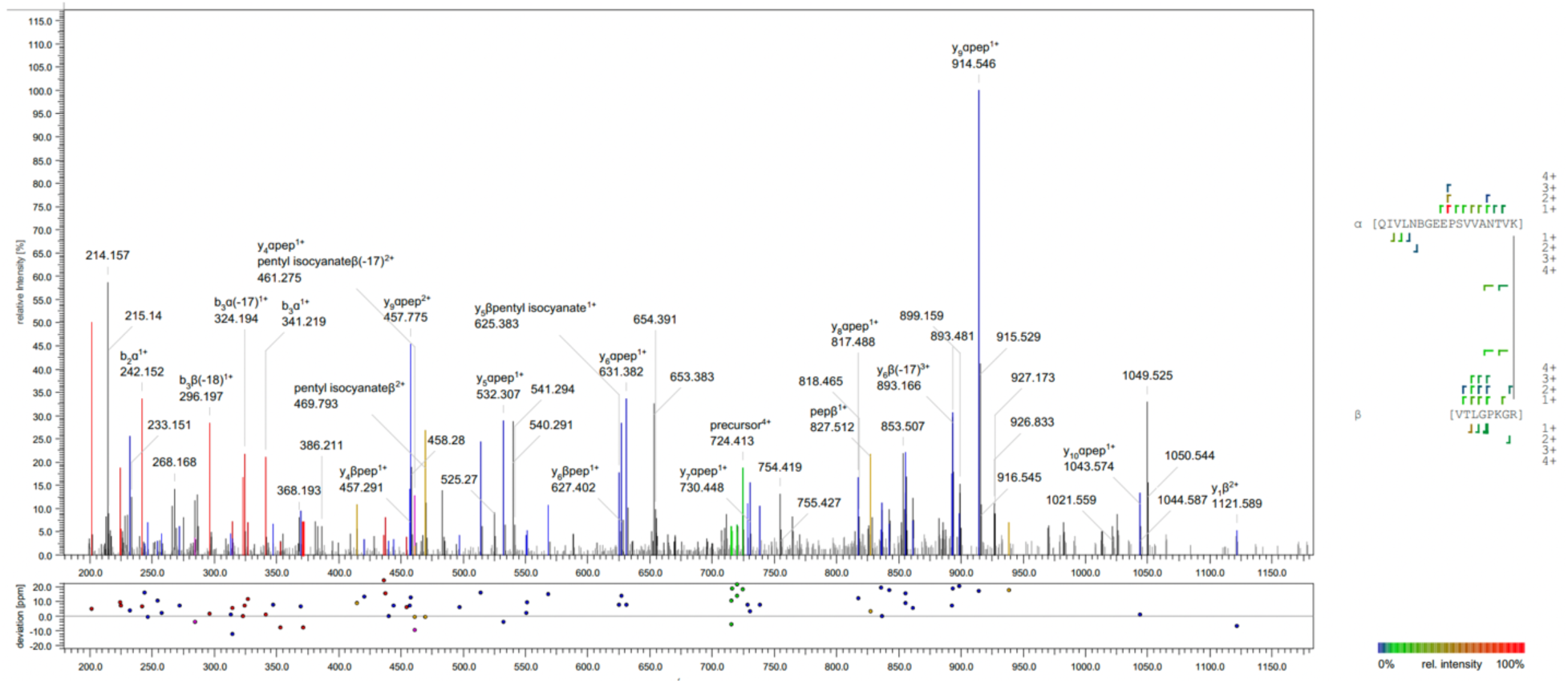
Representative spectrum from the DizSPC lysate experiment predigested with Lys-C. This spectrum shows an intraprotein crosslink in GroL between the ɑ-peptide [QIVLNBGEEPSVVANTVK] crosslinked at position K18 to the β-peptide, [VTLGPKGR], crosslinked at position R8.

**Figure S17.**
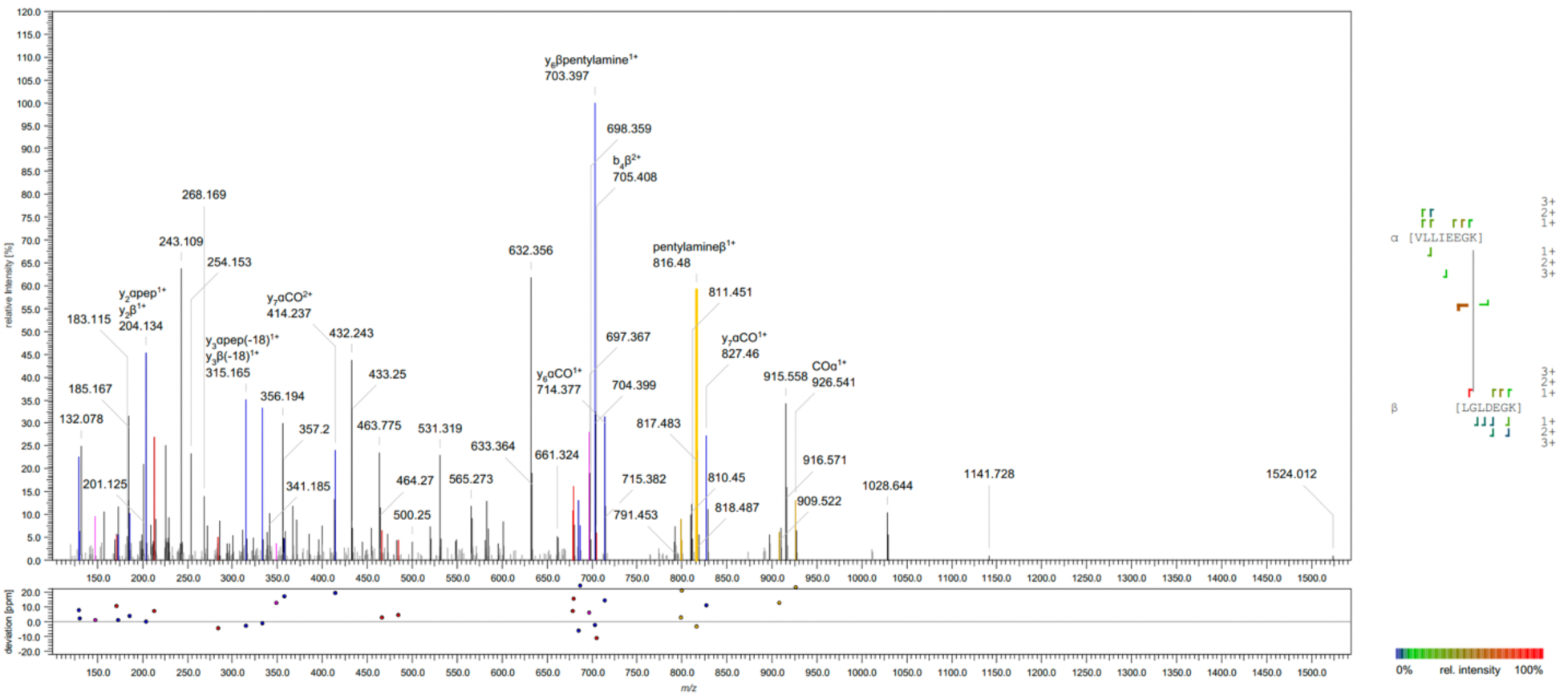
Representative spectrum from the DizSPC lysate experiment without Lys-C. This spectrum shows an interprotein crosslink between the ModE ɑ-peptide [VLLIEEGK] crosslinked at position K8 to the SsuB β-peptide, [LGLDEGK], crosslinked at position G2.

**Figure S18.**
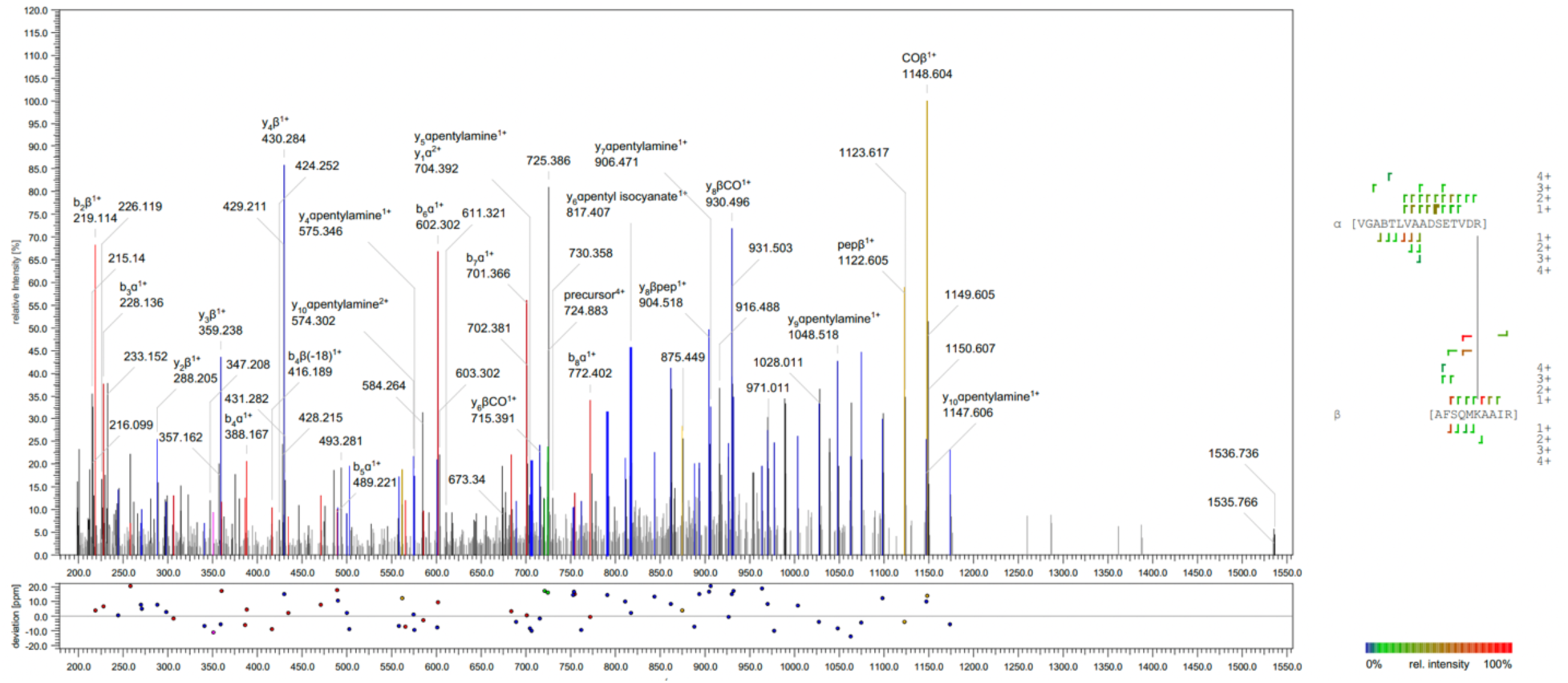
Representative spectrum from the DizSPC lysate experiment without Lys-C. This spectrum shows an intraprotein crosslink in aspC between the ɑ-peptide [VGABTLVAADSETVDR] crosslinked at position R16 to the β-peptide, [AFSQMKAAIR], crosslinked at position K6.

**Figure S19.**
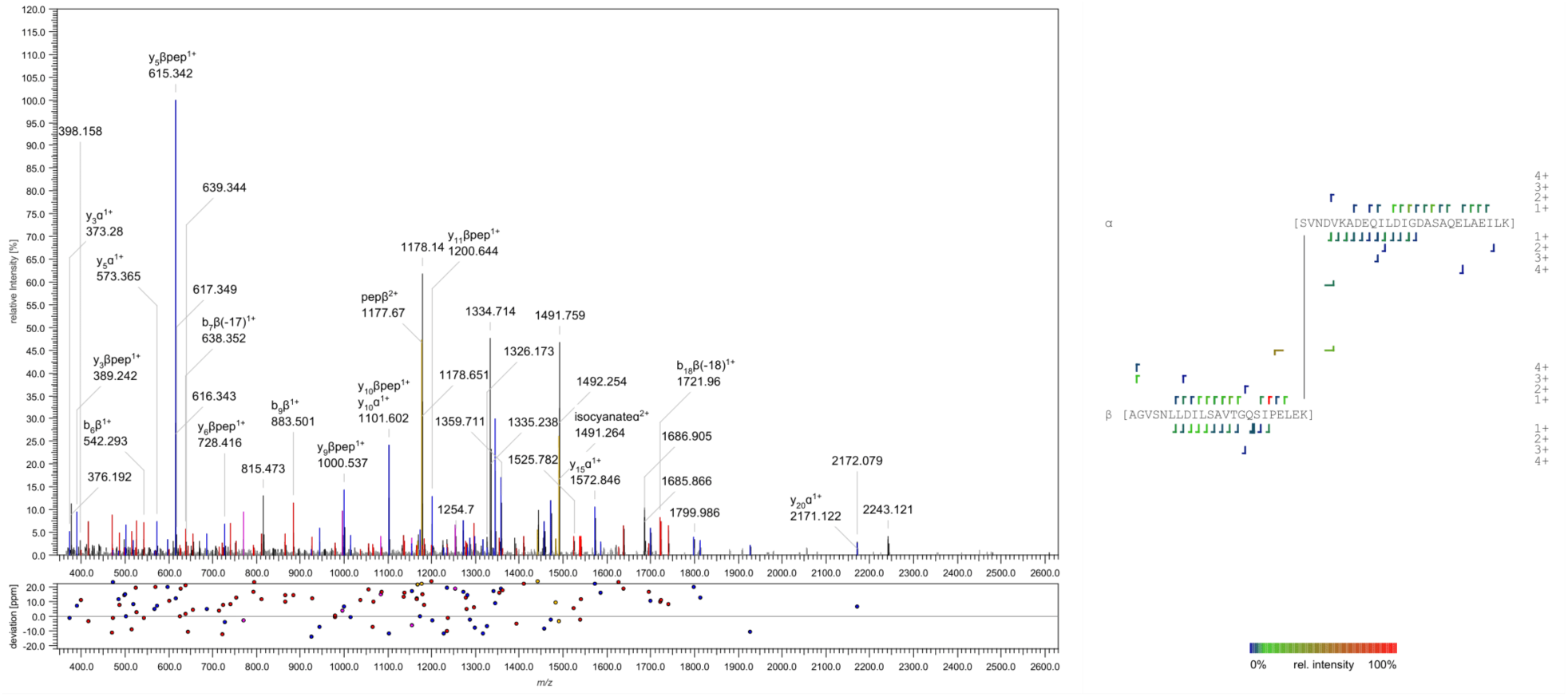
Representative spectrum from the in-cell DizSEC experiment. This spectrum shows an interprotein crosslink between the TrpS ɑ-peptide, [SVNDVKADEQILDIGDASAQELAEILK], crosslinked at position S1 to the Pgk β-peptide, [AGVSNLLDILSAVTGQSIPELEK], crosslinked at position K23.

**Figure S20:**
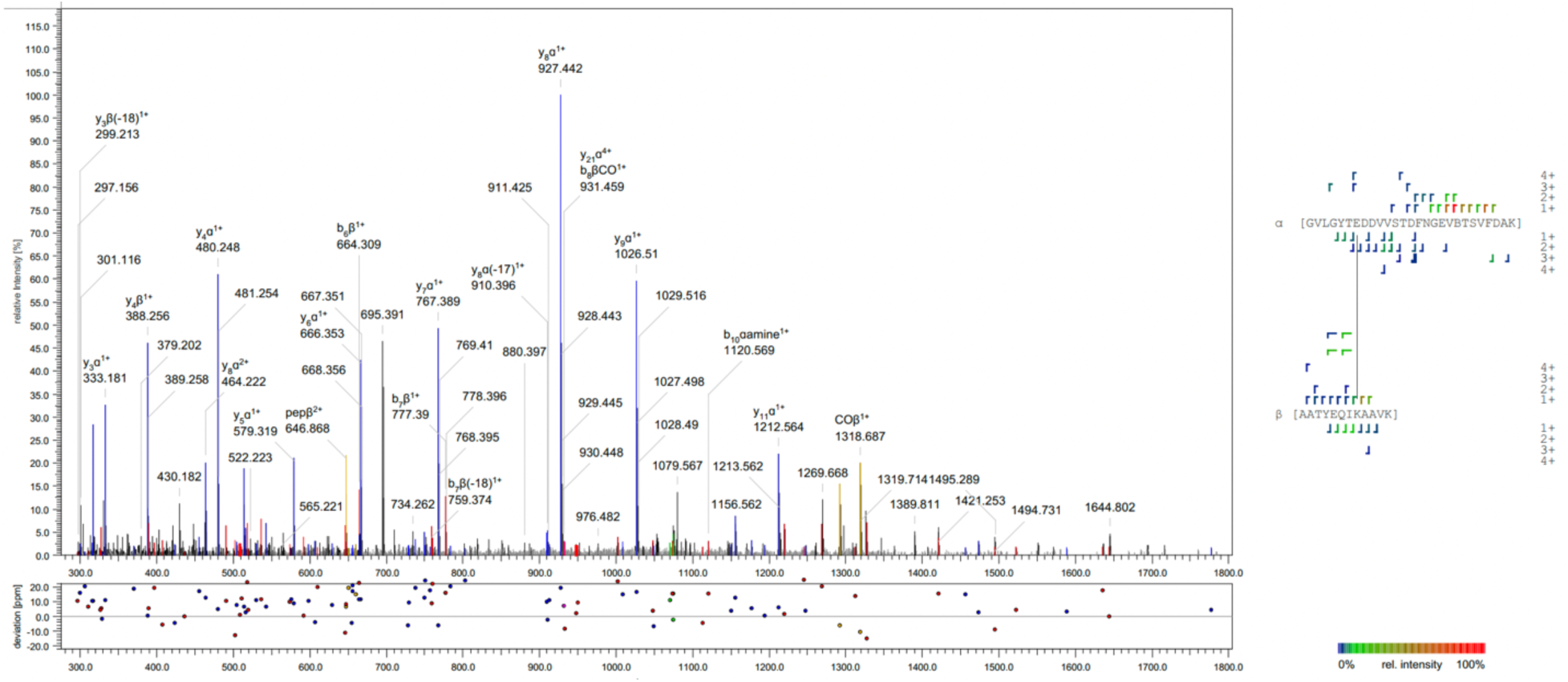
Representative spectrum from the in-cell DizSEC experiment. This spectrum shows an intraprotein crosslink in GapA between the ɑ-peptide [GVLGYTEDDVVSTDFNGEVBTSV-FDAK] crosslinked at position E7 to the β-peptide, [AATYEQIKAAVK], crosslinked at position K8.

**Figure S21.**
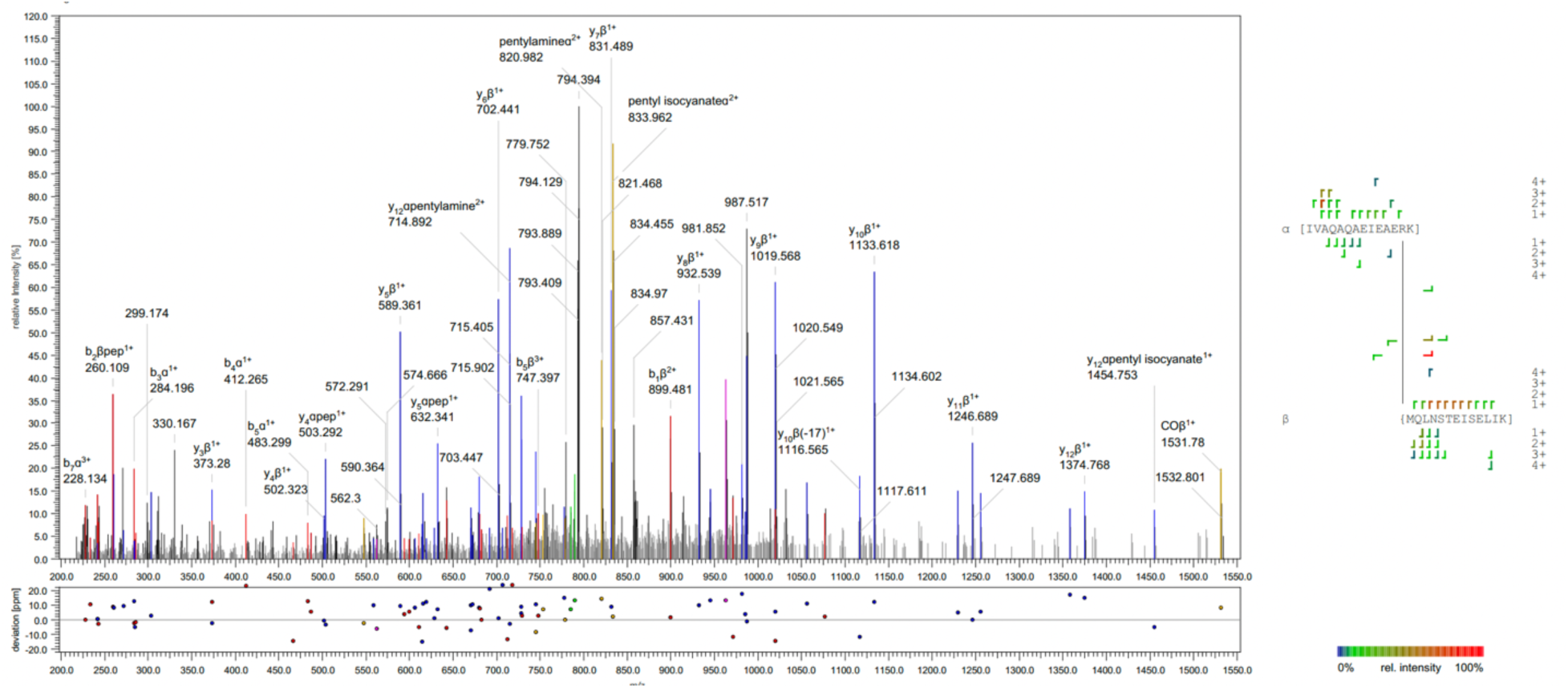
Representative spectrum from the in-cell DizSPC experiment. This spectrum shows an interprotein crosslink between the AtpA ɑ-peptide, [IVAQAQAEIEAERK], crosslinked at position R13 to the AtpF β-peptide, {MQLNSTEISELIK], crosslinked at N-terminus.

**Figure S22.**
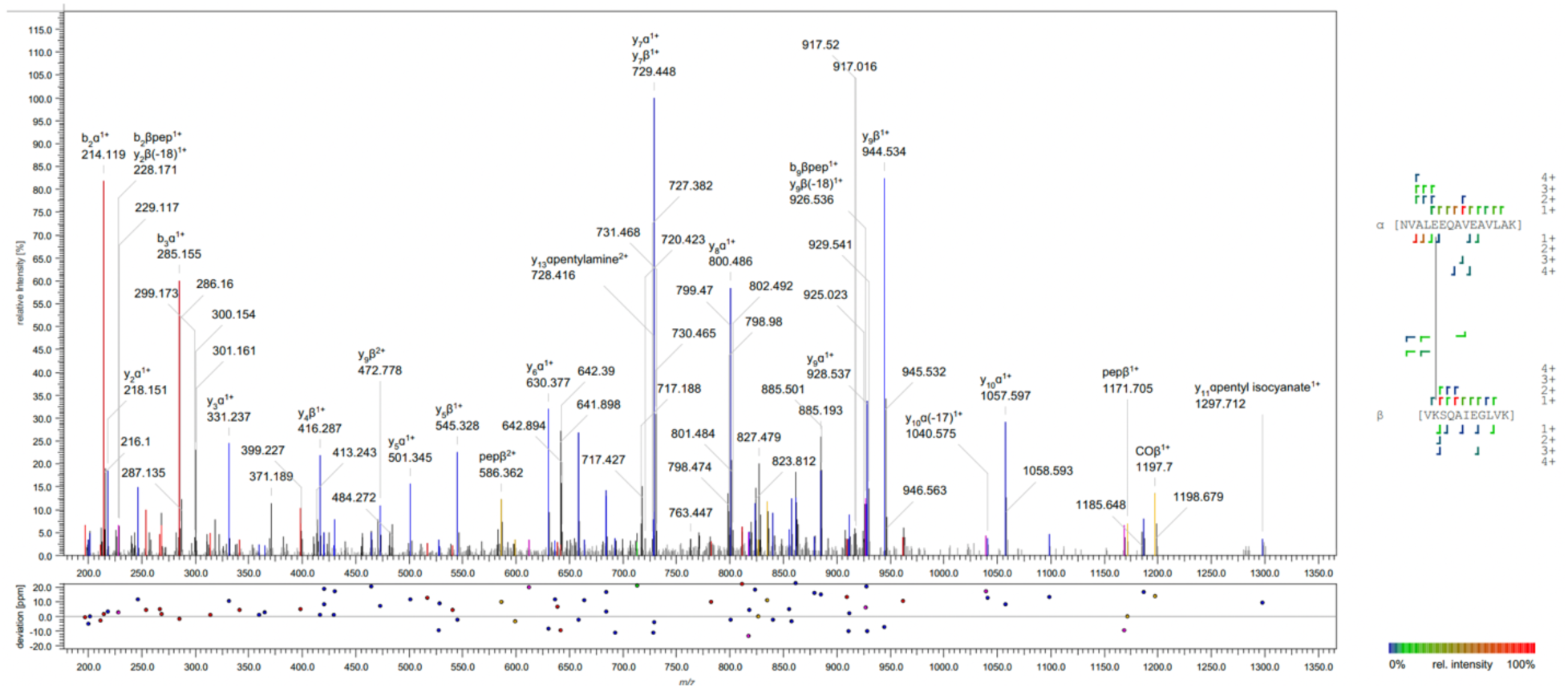
Representative spectrum from the in-cell DizSPC experiment. This spectrum shows an intraprotein crosslink in Tig between the ɑ-peptide [NVALEEQAVEAVLAK] crosslinked at position E5 to the β-peptide, [VKSQAIEGLVK], crosslinked at position K2.

### Section 4. Supplementary Tables

**Table S1.** Summary of DizSEC crosslinks on lysozyme with varied experimental and crosslinking conditions.

| DizSEC Conc. | UV% | LysC | Collisional Energy (%) | # XSMs<sup>a</sup> | XSM type |  |  | # Unique crosslinks<sup>b</sup> | Unique crosslink type |  |
| --- | --- | --- | --- | --- | --- | --- | --- | --- | --- | --- |
|  |  |  |  |  | Intra-peptidal | Dead-Ends | Inter-peptidal |  | Intra-protein | Homeotypic |
| - | 80%, 5 sec | + | 29, 32, 35 | 0 | 0 | 0 | 0 | 0 | 0 | 0 |
| 1 mM | - | + | 29, 32, 35 | 55 | 41 | 10 | 4 | 4 | 4 | 0 |
| 1 mM | 80%, 5 sec | - | 29, 32, 35 | 443 | 214 | 152 | 77 | 33 | 27 | 6 |
| 1 mM | 80%, 5 sec | + | 29, 32, 35 | 316 | 142 | 136 | 38 | 23 | 18 | 5 |
| 1 mM | 20%, 5 sec | + | 29, 32, 35 | 403 | 192 | 139 | 72 | 27 | 21 | 6 |
| 10 mM | 80%, 5 sec | + | 29, 32, 35 | 368 | 186 | 146 | 36 | 22 | 14 | 8 |
| 1 mM | 80%, 5 sec | + | 22, 25, 28 | 361 | 188 | 138 | 35 | 14 | 12 | 2 |
| 1 mM Sulfo-SDA | 80%, 5 sec | + | 29, 32, 35 | 348 | 196 | 80 | 72 | 22 | 19 | 3 |

<sup>a</sup> #XSMs include intrapeptidal, dead-end, and interpeptidal XSM types.
<sup>b</sup> #Unique crosslinks include interprotein, intraprotein, and homeotypic crosslink types.

**Table S2.** Summary of DizSPC crosslinks on lysozyme with varied experimental and crosslinking conditions.^a^

| DizSPC Conc. | UV% | LysC | DMAP | # XSMs<sup>b</sup> | XSM type |  |  | # Unique crosslinks<sup>c</sup> | Unique crosslink type |  |
| --- | --- | --- | --- | --- | --- | --- | --- | --- | --- | --- |
|  |  |  |  |  | Intra-peptidal | Dead-end | Inter-peptidal |  | Intra-protein | Homeotypic |
| - | 80%, 5 sec | + | - | 5 | 2 | 3 | 0 | 0 | 0 | 0 |
| 10 mM | - | + | - | 25 | 17 | 8 | 0 | 0 | 0 | 0 |
| 10 mM | 80%, 5 sec | + | - | 186 | 125 | 59 | 2 | 2 | 2 | 0 |
| 10 mM | 80%, 5 sec | + | 1 mM | 193 | 123 | 65 | 5 | 4 | 4 | 0 |
| 10 mM | 20%, 5 sec | - | - | 0 | 0 | 0 | 0 | 0 | 0 | 0 |
| 10 mM | 20%, 5 sec | + | - | 10 | 9 | 1 | 0 | 0 | 0 | 0 |
| 50 mM | 80%, 5 sec | + | - | 69 | 47 | 21 | 1 | 1 | 1 | 0 |

<sup>a</sup> All samples were fragmented with stepped normalized collisional energies of 29, 32, 35%, and used an isolation window of 1.0 *m/z*.
<sup>b</sup> #XSMs include intrapeptidal, dead-end, and interpeptidal XSM types.
<sup>c</sup> #Unique crosslinks include intraprotein and homeotypic crosslink types.

**Table S3.** Summary of DizSEC crosslinks from preliminary screening of enriched *E. coli* extracts.^a^

| SEC fraction | DMAP | Collisional Energy (%) | Isolation window | # XSMs<sup>b</sup> | XSM type |  |  | # Unique crosslinks<sup>c</sup> | Unique crosslink type |  |  |
| --- | --- | --- | --- | --- | --- | --- | --- | --- | --- | --- | --- |
|  |  |  |  |  | Intra-peptidal | Dead-end | Inter-peptidal |  | Inter-protein | Intra-protein | Homeotypic |
| 8 | - | 29, 32, 35 | 2.0 | 280 | 230 | 6 | 44 | 43 | 34 | 9 | 0 |
| 13 | - | 25, 28, 31 | 2.0 | 442 | 388 | 28 | 26 | 26 | 14 | 10 | 2 |
| 13 | - | 29, 32, 35 | 1.0 | 452 | 375 | 33 | 44 | 44 | 39 | 4 | 1 |
| 13 | - | 29, 32, 35 | 1.4 | 203 | 169 | 11 | 23 | 23 | 15 | 5 | 3 |
| 13 | - | 29, 32, 35 | 2.0 | 443 | 388 | 28 | 27 | 27 | 15 | 10 | 2 |
| 13 | - | 31, 34, 37 | 2.0 | 202 | 169 | 11 | 22 | 22 | 16 | 5 | 1 |
| 8 | + | 29, 32, 35 | 2.0 | 72 | 39 | 0 | 33 | 33 | 30 | 3 | 0 |
| 14 | + | 29, 32, 35 | 2.0 | 156 | 126 | 7 | 23 | 23 | 20 | 3 | 0 |

<sup>a</sup> All samples were exposed at a UV power of 80% for 5 s.<sup>b</sup> #XSMs include intrapeptidal, dead-end, and interpeptidal XSM types.<sup>c</sup> #Unique crosslinks include intraprotein and homeotypic crosslink types.

**Table S4.** Summary of DizSPC crosslinks from preliminary screening of enriched *E. coli* extracts.^a^

| SEC fraction | DMAP | Collisional Energy (%) | Isolation window | # XSMs<sup>b</sup> | XSM type |  |  | # Unique crosslinks<sup>c</sup> | Unique crosslink type |  |  |
| --- | --- | --- | --- | --- | --- | --- | --- | --- | --- | --- | --- |
|  |  |  |  |  | Intra-peptidal | Dead-end | Inter-peptidal |  | Inter-protein | Intra-protein | Homeotypic |
| 14 | + | 22, 25, 28 | 1.0 | 18 | 8 | 5 | 5 | 5 | 2 | 3 | 0 |
| 14 | + | 25, 28, 31 | 1.0 | 19 | 5 | 7 | 7 | 7 | 5 | 2 | 0 |
| 14 | + | 29, 32, 35 | 1.0 | 0 | 0 | 0 | 0 | 0 | 0 | 0 | 0 |
| 13 | + | 22, 25, 28 | 1.0 | 1486 | 1159 | 294 | 33 | 33 | 13 | 17 | 3 |
| 13 | + | 29, 32, 35 | 1.0 | 1574 | 1261 | 220 | 93 | 92 | 68 | 21 | 3 |

<sup>a</sup> All samples were exposed at a UV power of 80% for 5 s.<sup>b</sup> #XSMs include intrapeptidal, dead-end, and interpeptidal XSM types.<sup>c</sup> #Unique crosslinks include intraprotein and homeotypic crosslink types.

**Table S5.** Summary of DizSEC crosslinks from enriched *E. coli* extracts pre-digested by Lys-C.^a^

| SEC fraction | #XSMs<sup>b</sup> | XSM type |  |  | #Unique crosslinks<sup>c</sup> | Unique crosslink type |  |  |
| --- | --- | --- | --- | --- | --- | --- | --- | --- |
|  |  | Intra-peptidal | Dead-end | Inter-peptidal |  | Interproteins | Intra-peptidal | Dead-end |
| 4 | 67 | 46 | 18 | 3 | 3 | 3 | 0 | 0 |
| 5 | 29 | 18 | 7 | 4 | 4 | 4 | 0 | 0 |
| 6 | 72 | 44 | 17 | 11 | 11 | 11 | 0 | 0 |
| 7 | 373 | 234 | 82 | 57 | 57 | 57 | 0 | 0 |
| 8 | 269 | 191 | 64 | 16 | 14 | 12 | 2 | 0 |
| 9 | 508 | 354 | 140 | 16 | 14 | 12 | 1 | 1 |
| 10 | 434 | 320 | 94 | 25 | 20 | 15 | 1 | 4 |
| 11 | 478 | 325 | 121 | 34 | 32 | 30 | 1 | 1 |
| 14 | 364 | 257 | 81 | 28 | 26 | 24 | 0 | 2 |
| 15 | 482 | 353 | 114 | 21 | 15 | 9 | 1 | 5 |
| 16 | 298 | 224 | 49 | 28 | 25 | 22 | 3 | 0 |
| 17 | 217 | 89 | 95 | 33 | 33 | 33 | 0 | 0 |
| 18 | 97 | 49 | 34 | 15 | 14 | 13 | 0 | 1 |
| 19 | 61 | 34 | 18 | 9 | 9 | 9 | 0 | 0 |
| 20 | 78 | 61 | 13 | 4 | 4 | 4 | 0 | 0 |

<sup>a</sup> All samples were fragmented with stepped normalized collisional energies of 25, 28, 31%, used an isolation window of 1.0 *m/z*, and were treated with 1 mM DMAP.
<sup>b</sup> #XSMs include intrapeptidal, dead-end, and interpeptidal XSM types.
<sup>c</sup> #Unique crosslinks include interprotein, intraprotein, and homeotypic crosslink types.

**Table S6.**
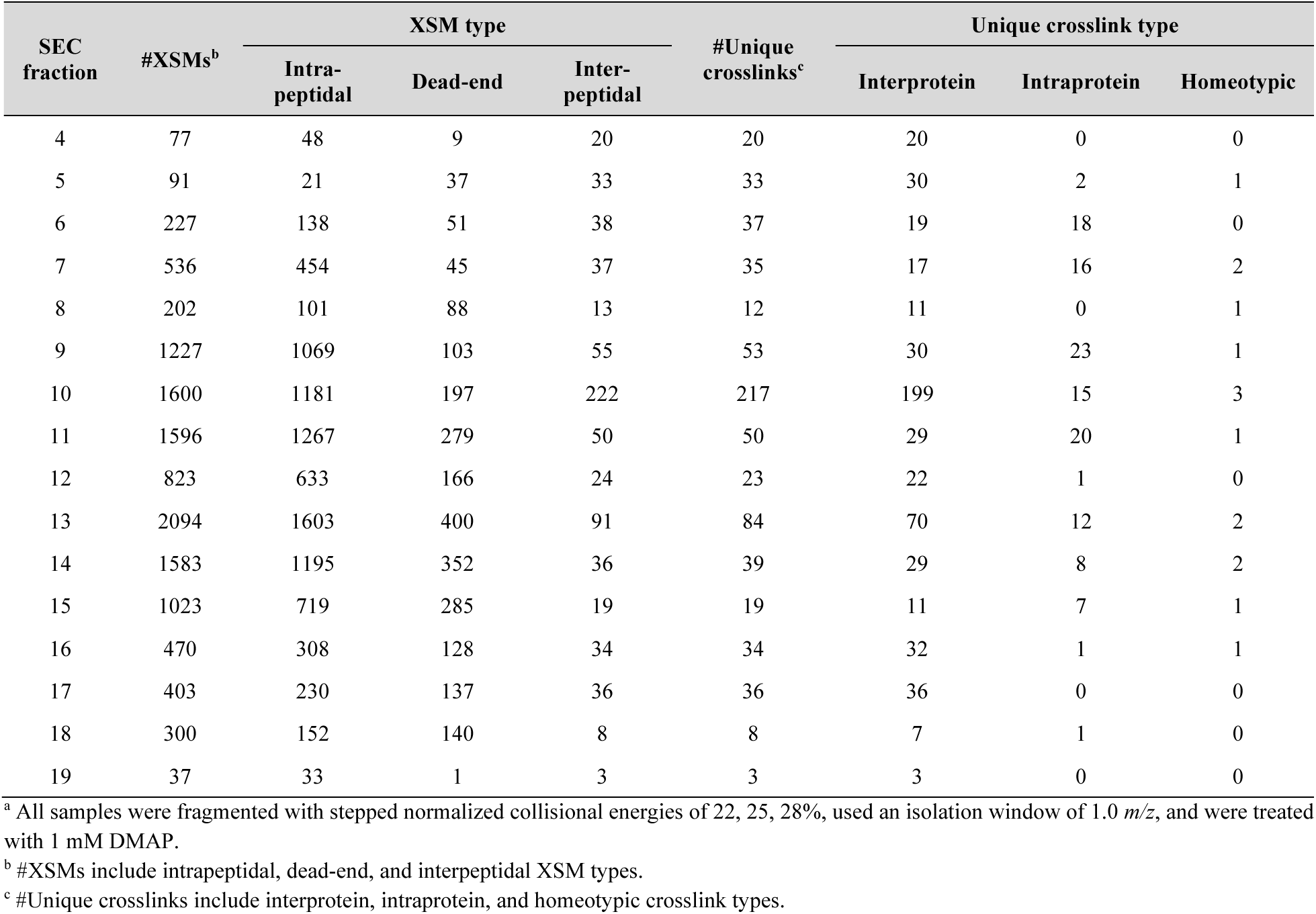
Summary of DizSEC crosslinks from enriched *E. coli* extracts without pre-digestion by Lys-C.^a^

**Table S7.** Summary of DizSPC crosslinks from enriched *E. coli* extracts pre-digested by Lys-C.^a^

| SEC fraction | #XSMs<sup>b</sup> | XSM type |  |  | #Unique crosslinks<sup>c</sup> | Unique crosslink type |  |  |
| --- | --- | --- | --- | --- | --- | --- | --- | --- |
|  |  | Intra-peptidal | Dead-end | Inter-peptidal |  | Interprotein | Intraprotein | Homeotypic |
| 4 | 33 | 30 | 2 | 1 | 1 | 1 | 0 | 0 |
| 5 | 63 | 23 | 6 | 34 | 34 | 33 | 0 | 1 |
| 6 | 140 | 79 | 27 | 34 | 34 | 34 | 0 | 0 |
| 7 | 68 | 45 | 3 | 20 | 18 | 16 | 0 | 2 |
| 8 | 531 | 429 | 61 | 41 | 35 | 17 | 13 | 5 |
| 9 | 545 | 354 | 155 | 36 | 35 | 19 | 13 | 3 |
| 10 | 610 | 479 | 92 | 39 | 38 | 22 | 15 | 1 |
| 11 | 676 | 452 | 195 | 29 | 28 | 9 | 18 | 1 |
| 12 | 722 | 491 | 124 | 107 | 107 | 92 | 15 | 0 |
| 13 | 1030 | 810 | 192 | 28 | 28 | 12 | 12 | 4 |
| 14 | 868 | 583 | 187 | 98 | 98 | 90 | 7 | 1 |
| 15 | 531 | 411 | 104 | 16 | 16 | 10 | 5 | 1 |
| 16 | 208 | 122 | 74 | 12 | 12 | 11 | 1 | 0 |
| 17 | 253 | 119 | 118 | 16 | 16 | 16 | 0 | 0 |
| 18 | 251 | 77 | 165 | 9 | 9 | 9 | 0 | 0 |
| 19 | 42 | 38 | 2 | 2 | 2 | 2 | 0 | 0 |

<sup>a</sup> All samples were fragmented with stepped normalized collisional energies of 22, 25, 28%, used an isolation window of 1.0 *m/z*, and were treated with 1 mM DMAP.
<sup>b</sup> #XSMs include intrapeptidal, dead-end, and interpeptidal XSM types.
<sup>c</sup> #Unique crosslinks include interprotein, intraprotein, and homeotypic crosslink types.

**Table S8.**
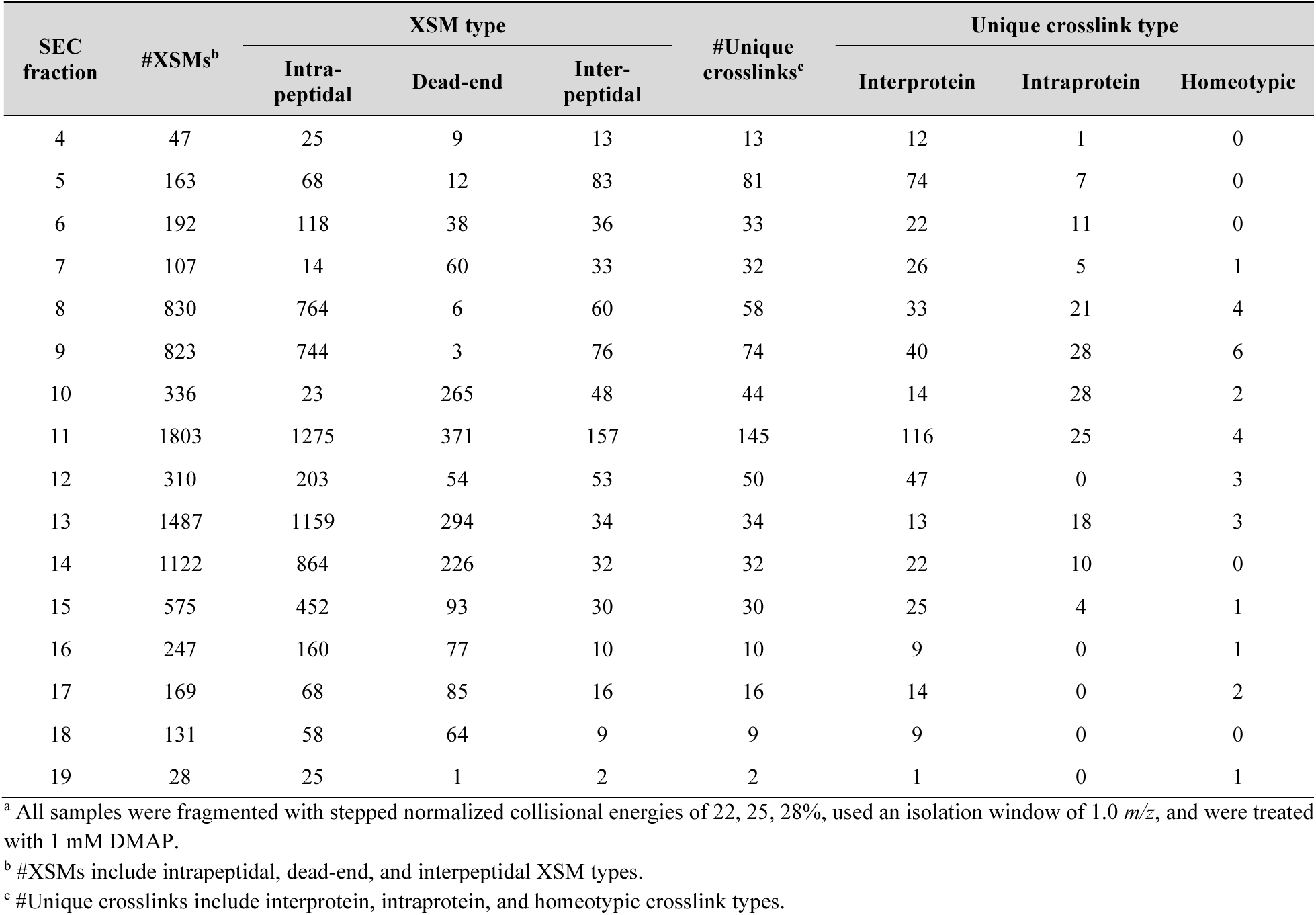
Summary of DizSPC crosslinks from enriched *E. coli* extracts without pre-digestion by Lys-C.^a^

**Table S9.** Summary of DizSEC crosslinks from enriched in-cell samples pre-digested by Lys-C.^a^

| SEC fraction | #XSMs<sup>b</sup> | XSM type |  |  | #Unique crosslinks<sup>c</sup> | Unique crosslink type |  |  |
| --- | --- | --- | --- | --- | --- | --- | --- | --- |
|  |  | Intra-peptidal | Dead-end | Inter-peptidal |  | Interprotein | Intraprotein | Homeotypic |
| 4 | 43 | 3 | 19 | 21 | 19 | 16 | 3 | 0 |
| 5 | 170 | 51 | 72 | 47 | 47 | 33 | 12 | 2 |
| 6 | 226 | 100 | 59 | 67 | 59 | 12 | 46 | 1 |
| 7 | 501 | 295 | 82 | 124 | 114 | 43 | 69 | 2 |
| 8 | 1005 | 781 | 125 | 99 | 95 | 27 | 67 | 1 |
| 9 | 1778 | 1363 | 269 | 146 | 141 | 26 | 109 | 6 |
| 10 | 2214 | 1738 | 314 | 162 | 160 | 55 | 101 | 4 |
| 11 | 2070 | 1726 | 180 | 164 | 159 | 69 | 87 | 3 |
| 12 | 2431 | 2044 | 267 | 120 | 120 | 40 | 78 | 2 |
| 13 | 2847 | 2297 | 410 | 140 | 137 | 34 | 98 | 5 |
| 14 | 2780 | 2222 | 435 | 123 | 123 | 64 | 57 | 2 |
| 15 | 2607 | 1981 | 530 | 96 | 96 | 51 | 44 | 1 |
| 16 | 2306 | 1805 | 406 | 95 | 94 | 57 | 35 | 2 |
| 17 | 1850 | 1552 | 253 | 45 | 45 | 13 | 30 | 2 |
| 18 | 1328 | 1018 | 253 | 57 | 56 | 38 | 17 | 1 |
| 19 | 409 | 273 | 115 | 21 | 21 | 18 | 2 | 1 |

<sup>a</sup> All samples were fragmented with stepped normalized collisional energies of 22, 25, 28% and used an isolation window of 1.0 *m/z*.<sup>b</sup> #XSMs include intrapeptidal, dead-end, and interpeptidal XSM types.<sup>c</sup> #Unique crosslinks include interprotein, intraprotein, and homeotypic crosslink types.

**Table S10.** Summary of DizSPC crosslinks from enriched in-cell samples pre-digested by Lys-C.^a^

| SEC fraction | #XSMs<sup>b</sup> | XSM type |  |  | #Unique crosslinks<sup>c</sup> | Unique crosslink type |  |  |
| --- | --- | --- | --- | --- | --- | --- | --- | --- |
|  |  | Intra-peptidal | Dead-end | Inter-peptidal |  | Interprotein | Intraprotein | Homeotypic |
| 4 | 190 | 75 | 63 | 52 | 54 | 48 | 3 | 1 |
| 5 | 258 | 106 | 93 | 59 | 55 | 29 | 24 | 2 |
| 6 | 309 | 132 | 70 | 107 | 91 | 24 | 65 | 2 |
| 7 | 718 | 522 | 91 | 105 | 99 | 24 | 72 | 3 |
| 8 | 1407 | 1036 | 273 | 98 | 91 | 14 | 75 | 2 |
| 9 | 1854 | 1477 | 237 | 140 | 138 | 58 | 74 | 6 |
| 10 | 2043 | 1654 | 264 | 125 | 121 | 21 | 93 | 7 |
| 11 | 2184 | 1757 | 296 | 131 | 126 | 34 | 90 | 2 |
| 13 | 2535 | 2095 | 351 | 89 | 87 | 15 | 69 | 3 |
| 14 | 2476 | 2017 | 382 | 77 | 74 | 24 | 49 | 1 |
| 15 | 2100 | 1696 | 333 | 71 | 67 | 38 | 27 | 2 |
| 16 | 1835 | 1299 | 476 | 60 | 58 | 39 | 17 | 2 |
| 17 | 1328 | 911 | 358 | 59 | 57 | 34 | 23 | 0 |
| 18 | 985 | 676 | 276 | 33 | 31 | 13 | 16 | 2 |
| 19 | 330 | 166 | 142 | 22 | 22 | 19 | 1 | 2 |

<sup>a</sup> All samples were fragmented with stepped normalized collisional energies of 22, 25, 28% and used an isolation window of 1.0 *m/z*.
<sup>b</sup> #XSMs include intrapeptidal, dead-end, and interpeptidal XSM types.
<sup>c</sup> #Unique crosslinks include interprotein, intraprotein, and homeotypic crosslink types.

## Notes

### Competing Interest Statement

The authors have declared no competing interest.

## REFERENCES

(1) Iacobucci, C.; Götze, M.; Sinz, A. Cross-Linking/Mass Spectrometry to Get a Closer View on Protein Interaction Networks. Curr Opin Biotechnol 2020, 63, 48–53.

(2) Mishra, P. K.; Yoo, C.; Hong, E.; Rhee, H. W. Photo-crosslinking: An Emerging Chemical Tool for Investigating Molecular Networks in Live Cells. ChemBioChem 2020, 21 (7), 924–932.

(3) Lenz, S.; Sinn, L. R.; O’Reilly, F. J.; Fischer, L.; Wegner, F.; Rappsilber, J. Reliable Identification of Protein-Protein Interactions by Crosslinking Mass Spectrometry. Nat Commun 2021, 12 (1), 1–11.

(4) Wheat, A.; Yu, C.; Wang, X.; Burke, A. M.; Chemmama, I. E.; Kaake, R. M.; Baker, P.; Rychnovsky, S. D.; Yang, J.; Huang, L. Protein Interaction Landscapes Revealed by Advanced in Vivo Cross-Linking–Mass Spectrometry. Proceedings of the National Academy of Sciences 2021, 118 (32), e2023360118.

(5) O’Reilly, F. J.; Xue, L.; Graziadei, A.; Sinn, L.; Lenz, S.; Tegunov, D.; Blötz, C.; Singh, N.; Hagen, W. J. H.; Cramer, P. In-Cell Architecture of an Actively Transcribing-Translating Expressome. Science (1979) 2020, 369 (6503), 554–557.

(6) Leitner, A.; Walzthoeni, T.; Kahraman, A.; Herzog, F.; Rinner, O.; Beck, M.; Aebersold, R. Probing Native Protein Structures by Chemical Cross-Linking, Mass Spectrometry, and Bioinformatics. Molecular & Cellular Proteomics 2010, 9 (8), 1634–1649.

(7) Sinz, A. Chemical Cross-linking and Mass Spectrometry for Mapping Three-dimensional Structures of Proteins and Protein Complexes. Journal of mass spectrometry 2003, 38 (12), 1225–1237.

(8) Trakselis, M. A.; Alley, S. C.; Ishmael, F. T. Identification and Mapping of Protein− Protein Interactions by a Combination of Cross-Linking, Cleavage, and Proteomics. Bioconjug Chem 2005, 16 (4), 741–750.

(9) Bishop, A. C.; Buzko, O.; Shokat, K. M. Magic Bullets for Protein Kinases. Trends Cell Biol 2001, 11 (4), 167–172.

(10) Petrotchenko, E. v; Borchers, C. H. Crosslinking Combined with Mass Spectrometry for Structural Proteomics. Mass Spectrom Rev 2010, 29 (6), 862–876.

(11) Tang, X.; Bruce, J. E. A New Cross-Linking Strategy: Protein Interaction Reporter (PIR) Technology for Protein–Protein Interaction Studies. Mol Biosyst 2010, 6 (6), 939–947.

(12) Liu, F.; Rijkers, D. T. S.; Post, H.; Heck, A. J. R. Proteome-Wide Profiling of Protein Assemblies by Cross-Linking Mass Spectrometry. Nat Methods 2015, 12 (12), 1179–1184.

(13) Arora, B.; Tandon, R.; Attri, P.; Bhatia, R. Chemical Crosslinking: Role in Protein and Peptide Science. Curr Protein Pept Sci 2017, 18 (9), 946–955.

(14) Matzinger, M.; Mechtler, K. Cleavable Cross-Linkers and Mass Spectrometry for the Ultimate Task of Profiling Protein–Protein Interaction Networks in Vivo. J Proteome Res 2020, 20 (1), 78–93.

(15) O’Reilly, F. J.; Rappsilber, J. Cross-Linking Mass Spectrometry: Methods and Applications in Structural, Molecular and Systems Biology. Nat Struct Mol Biol 2018, 25 (11), 1000–1008.

(16) Götze, M.; Iacobucci, C.; Ihling, C. H.; Sinz, A. A Simple Cross-Linking/Mass Spectrometry Workflow for Studying System-Wide Protein Interactions. Anal Chem 2019, 91 (15), 10236–10244. https://doi.org/10.1021/acs.analchem.9b02372.

(17) Xue, L.; Lenz, S.; Zimmermann-Kogadeeva, M.; Tegunov, D.; Cramer, P.; Bork, P.; Rappsilber, J.; Mahamid, J. Visualizing Translation Dynamics at Atomic Detail inside a Bacterial Cell. bioRxiv 2021.

(18) Schweppe, D. K.; Chavez, J. D.; Lee, C. F.; Caudal, A.; Kruse, S. E.; Stuppard, R.; Marcinek, D. J.; Shadel, G. S.; Tian, R.; Bruce, J. E. Mitochondrial Protein Interactome Elucidated by Chemical Cross-Linking Mass Spectrometry. Proceedings of the National Academy of Sciences 2017, 114 (7), 1732–1737.

(19) Chavez, J. D.; Lee, C. F.; Caudal, A.; Keller, A.; Tian, R.; Bruce, J. E. Chemical Crosslinking Mass Spectrometry Analysis of Protein Conformations and Supercomplexes in Heart Tissue. Cell Syst 2018, 6 (1), 136–141.

(20) Klykov, O.; Steigenberger, B.; Pektaş, S.; Fasci, D.; Heck, A. J. R.; Scheltema, R. A. Efficient and Robust Proteome-Wide Approaches for Cross-Linking Mass Spectrometry. Nat Protoc 2018, 13 (12), 2964–2990.

(21) Kaake, R. M.; Wang, X.; Burke, A.; Yu, C.; Kandur, W.; Yang, Y.; Novtisky, E. J.; Second, T.; Duan, J.; Kao, A. A New in Vivo Cross-Linking Mass Spectrometry Platform to Define Protein–Protein Interactions in Living Cells. Molecular & Cellular Proteomics 2014, 13 (12), 3533–3543.

(22) Moss, R. A. Diazirines: Carbene Precursors Par Excellence. Acc Chem Res 2006, 39 (4), 267–272.

(23) Das, J. Aliphatic Diazirines as Photoaffinity Probes for Proteins: Recent Developments. Chem Rev 2011, 111 (8), 4405–4417.

(24) Deng, H.; Lei, Q.; Wu, Y.; He, Y.; Li, W. Activity-Based Protein Profiling: Recent Advances in Medicinal Chemistry. Eur J Med Chem 2020, 191, 112151.

(25) Hu, W.; Yuan, Y.; Wang, C.-H.; Tian, H.-T.; Guo, A.-D.; Nie, H.-J.; Hu, H.; Tan, M.; Tang, Z.; Chen, X.-H. Genetically Encoded Residue-Selective Photo-Crosslinker to Capture Protein-Protein Interactions in Living Cells. Chem 2019, 5 (11), 2955–2968.

(26) Dubinsky, L.; Krom, B. P.; Meijler, M. M. Diazirine Based Photoaffinity Labeling. Bioorg Med Chem 2012, 20 (2), 554–570.

(27) Zhuo, F.; Guo, Q.; Zheng, Y.; Liu, T.; Yang, Z.; Xu, Q.; Jiang, Y.; Liu, D.; Zeng, K.; Tu, P. Photoaffinity Labeling-Based Chemoproteomic Strategy Reveals RBBP4 as a Cellular Target of Protopanaxadiol against Colorectal Cancer Cells. ChemBioChem 2022, e202200038.

(28) West, A. v; Muncipinto, G.; Wu, H.-Y.; Huang, A. C.; Labenski, M. T.; Jones, L. H.; Woo, C. M. Labeling Preferences of Diazirines with Protein Biomolecules. J Am Chem Soc 2021, 143 (17), 6691–6700.

(29) Halloran, M. W.; Lumb, J. Recent Applications of Diazirines in Chemical Proteomics. Chemistry–A European Journal 2019, 25 (19), 4885–4898.

(30) Toscano, J. P.; Platz, M. S.; Nikolaev, V. Lifetimes of Simple Ketocarbenes. J Am Chem Soc 1995, 117 (16), 4712–4713.

(31) Yang, T.; Liu, Z.; Li, X. D. Developing Diazirine-Based Chemical Probes to Identify Histone Modification ‘Readers’ and ‘Erasers.’ Chem Sci 2015, 6 (2), 1011–1017.

(32) Yang, T.; Li, X.-M.; Bao, X.; Fung, Y. M. E.; Li, X. D. Photo-Lysine Captures Proteins That Bind Lysine Post-Translational Modifications. Nat Chem Biol 2016, 12 (2), 70–72.

(33) Yang, Y.; Song, H.; He, D.; Zhang, S.; Dai, S.; Lin, S.; Meng, R.; Wang, C.; Chen, P. R. Genetically Encoded Protein Photocrosslinker with a Transferable Mass Spectrometry-Identifiable Label. Nat Commun 2016, 7 (1), 1–10.

(34) Lin, S.; He, D.; Long, T.; Zhang, S.; Meng, R.; Chen, P. R. Genetically Encoded Cleavable Protein Photo-Cross-Linker. J Am Chem Soc 2014, 136 (34), 11860–11863.

(35) Kalkhof, S.; Sinz, A. Chances and Pitfalls of Chemical Cross-Linking with Amine-Reactive N-Hydroxysuccinimide Esters. Anal Bioanal Chem 2008, 392 (1), 305–312.

(36) Wang, J.; Kubicki, J.; Peng, H.; Platz, M. S. Influence of Solvent on Carbene Intersystem Crossing Rates. J Am Chem Soc 2008, 130 (20), 6604–6609.

(37) Blencowe, A.; Hayes, W. Development and Application of Diazirines in Biological and Synthetic Macromolecular Systems. Soft Matter 2005, 1 (3), 178–205.

(38) Belsom, A.; Rappsilber, J. Anatomy of a Crosslinker. Curr Opin Chem Biol 2021, 60, 39–46.

(39) Iacobucci, C.; Piotrowski, C.; Aebersold, R.; Amaral, B. C.; Andrews, P.; Bernfur, K.; Borchers, C.; Brodie, N. I.; Bruce, J. E.; Cao, Y. First Community-Wide, Comparative Cross-Linking Mass Spectrometry Study. Anal Chem 2019, 91 (11), 6953–6961.

(40) Nury, C.; Redeker, V.; Dautrey, S.; Romieu, A.; van der Rest, G.; Renard, P.-Y.; Melki, R.; Chamot-Rooke, J. A Novel Bio-Orthogonal Cross-Linker for Improved Protein/Protein Interaction Analysis. Anal Chem 2015, 87 (3), 1853–1860.

(41) Mahamid, J.; Pfeffer, S.; Schaffer, M.; Villa, E.; Danev, R.; Kuhn Cuellar, L.; Förster, F.; Hyman, A. A.; Plitzko, J. M.; Baumeister, W. Visualizing the Molecular Sociology at the HeLa Cell Nuclear Periphery. Science (1979) 2016, 351 (6276), 969–972.

(42) Pfeffer, S.; Mahamid, J. Unravelling Molecular Complexity in Structural Cell Biology. Curr Opin Struct Biol 2018, 52, 111–118.

(43) Freer, R.; McKillop, A. Synthesis of Symmetrical and Unsymmetrical Ureas Using Unsymmetrical Diaryl Carbonates. Synth Commun 1996, 26 (2), 331–349.

(44) Höfle, G.; Steglich, W.; Vorbrüggen, H. 4-Dialkylaminopyridines as Highly Active Acylation Catalysts.[New Synthetic Method (25)]. Angewandte Chemie International Edition in English 1978, 17 (8), 569–583.

(45) Chambers, M. C.; Maclean, B.; Burke, R.; Amodei, D.; Ruderman, D. L.; Neumann, S.; Gatto, L.; Fischer, B.; Pratt, B.; Egertson, J. A Cross-Platform Toolkit for Mass Spectrometry and Proteomics. Nat Biotechnol 2012, 30 (10), 918–920.

(46) Götze, M.; Pettelkau, J.; Schaks, S.; Bosse, K.; Ihling, C. H.; Krauth, F.; Fritzsche, R.; Kühn, U.; Sinz, A. StavroX—a Software for Analyzing Crosslinked Products in Protein Interaction Studies. J Am Soc Mass Spectrom 2011, 23 (1), 76–87.

(47) Graham, M.; Combe, C.; Kolbowski, L.; Rappsilber, J. XiView: A Common Platform for the Downstream Analysis of Crosslinking Mass Spectrometry Data. BioRxiv 2019, 561829.

(48) Iacobucci, C.; Götze, M.; Ihling, C. H.; Piotrowski, C.; Arlt, C.; Schafer, M.; Hage, C.; Schmidt, R.; Sinz, A. A Cross-Linking/Mass Spectrometry Workflow Based on MS-Cleavable Cross-Liners and the MeroX Software for Studying Protein Structures and Protein-Protein Interactions. Nat. Protoc. 2018, 13, 2864–2889.

(49) Vellucci, D.; Kao, A.; Kaake, R. M.; Rychnovsky, S. D.; Huang, L. Selective Enrichment and Identification of Azide-Tagged Cross-Linked Peptides Using Chemical Ligation and Mass Spectrometry. J Am Soc Mass Spectrom 2010, 21 (8), 1432–1445.

(50) Fritzsche, R.; Ihling, C. H.; Götze, M.; Sinz, A. Optimizing the Enrichment of Cross-Linked Products for Mass Spectrometric Protein Analysis. Rapid Communications in Mass Spectrometry 2012, 26 (6), 653–658.

(51) Schmidt, R.; Sinz, A. Improved Single-Step Enrichment Methods of Cross-Linked Products for Protein Structure Analysis and Protein Interaction Mapping. Anal Bioanal Chem 2017, 409 (9), 2393–2400.

(52) Hulsen, T.; de Vlieg, J.; Alkema, W. BioVenn–a Web Application for the Comparison and Visualization of Biological Lists Using Area-Proportional Venn Diagrams. BMC Genomics 2008, 9 (1), 1–6.

(53) Keller, A.; Chavez, J. D.; Felt, K. C.; Bruce, J. E. Prediction of an Upper Limit for the Fraction of Interprotein Cross-Links in Large-Scale in Vivo Cross-Linking Studies. J Proteome Res 2019, 18 (8), 3077–3085.

(54) Dunkle, J. A.; Wang, L.; Feldman, M. B.; Pulk, A.; Chen, V. B.; Kapral, G. J.; Noeske, J.; Richardson, J. S.; Blanchard, S. C.; Cate, J. H. D. Structures of the Bacterial Ribosome in Classical and Hybrid States of TRNA Binding. Science (1979) 2011, 332 (6032), 981–984.

(55) Pulk, A.; Cate, J. H. D. Control of Ribosomal Subunit Rotation by Elongation Factor G. Science (1979) 2013, 340 (6140).

(56) Montesano-Roditis, L.; Glitz, D. G.; Traut, R. R.; Stewart, P. L. Cryo-Electron Microscopic Localization of Protein L7/L12 within the Escherichia Coli 70 S Ribosome by Difference Mapping and Nanogold Labeling. Journal of Biological Chemistry 2001, 276 (17), 14117–14123.

(57) Liljas, A.; Sanyal, S. The Enigmatic Ribosomal Stalk. Q Rev Biophys 2018, 51.

(58) Wendrich, T. M.; Blaha, G.; Wilson, D. N.; Marahiel, M. A.; Nierhaus, K. H. Dissection of the Mechanism for the Stringent Factor RelA. Mol Cell 2002, 10 (4), 779–788.

(59) Zhao, G.; Winkler, M. E. A Novel Alpha-Ketoglutarate Reductase Activity of the SerA-Encoded 3-Phosphoglycerate Dehydrogenase of Escherichia Coli K-12 and Its Possible Implications for Human 2-Hydroxyglutaric Aciduria. J Bacteriol 1996, 178 (1), 232–239.

(60) Stepp, L. R.; Bleile, D. M.; McRorie, D. K.; Pettit, F. H.; Reed, L. J. Use of Trypsin and Lipoamidase to Study the Role of Lipoic Acid Moieties in the Pyruvate and. Alpha.-Ketoglutarate Dehydrogenase Complexes of Escherichia Coli. Biochemistry 1981, 20 (16), 4555–4560.

(61) Guo, X.; Myasnikov, A. G.; Chen, J.; Crucifix, C.; Papai, G.; Takacs, M.; Schultz, P.; Weixlbaumer, A. Structural Basis for NusA Stabilized Transcriptional Pausing. Mol Cell 2018, 69 (5), 816–827.

(62) Hammel, M.; Amlanjyoti, D.; Reyes, F. E.; Chen, J.-H.; Parpana, R.; Tang, H. Y. H.; Larabell, C. A.; Tainer, J. A.; Adhya, S. HU Multimerization Shift Controls Nucleoid Compaction. Sci Adv 2016, 2 (7), e1600650–e1600650.

(63) Stelmashenko, O.; Lalo, U.; Yang, Y.; Bragg, L.; North, R. A.; Compan, V. Activation of Trimeric P2X2 Receptors by Fewer than Three ATP Molecules. Mol Pharmacol 2012, 82 (4), 760–766.

(64) Ghosh, P.; Ishihama, A.; Chatterji, D. Escherichia Coli RNA Polymerase Subunit ω and Its N-terminal Domain Bind Full-length Β′ to Facilitate Incorporation into the Α2β Subassembly. Eur J Biochem 2001, 268 (17), 4621–4627.

(65) Kar, S.; Edgar, R.; Adhya, S. Nucleoid Remodeling by an Altered HU Protein: Reorganization of the Transcription Program. Proceedings of the National Academy of Sciences 2005, 102 (45), 16397–16402.

(66) Calloni, G.; Chen, T.; Schermann, S. M.; Chang, H. C.; Genevaux, P.; Agostini, F.; Tartaglia, G. G.; Hayer-Hartl, M.; Hartl, F. U. DnaK Functions as a Central Hub in the E. Coli Chaperone Network. Cell Rep 1 (3): 251–264. 2012.

(67) Kerner, M. J.; Naylor, D. J.; Ishihama, Y.; Maier, T.; Chang, H.-C.; Stines, A. P.; Georgopoulos, C.; Frishman, D.; Hayer-Hartl, M.; Mann, M. Proteome-Wide Analysis of Chaperonin-Dependent Protein Folding in Escherichia Coli. Cell 2005, 122 (2), 209–220.

(68) Mayer, M. P.; Prodromou, C.; Frydman, J. The Hsp90 Mosaic: A Picture Emerges. Nat Struct Mol Biol 2009, 16 (1), 2–6.

(69) Chiosis, G.; Dickey, C. A.; Johnson, J. L. A Global View of Hsp90 Functions. Nat Struct Mol Biol 2013, 20 (1), 1–4.

(70) Marx, D. C.; Plummer, A. M.; Faustino, A. M.; Devlin, T.; Roskopf, M. A.; Leblanc, M. J.; Lessen, H. J.; Amann, B. T.; Fleming, P. J.; Krueger, S. SurA Is a Cryptically Grooved Chaperone That Expands Unfolded Outer Membrane Proteins. Proceedings of the National Academy of Sciences 2020, 117 (45), 28026–28035.

(71) Grudniak, A. M.; Pawlak, K.; Bartosik, K.; Wolska, K. I. Physiological Consequences of Mutations in the HtpG Heat Shock Gene of Escherichia Coli. Mutation Research/Fundamental and Molecular Mechanisms of Mutagenesis 2013, 745, 1–5.

(72) Karp, P. D.; Ong, W. K.; Paley, S.; Billington, R.; Caspi, R.; Fulcher, C.; Kothari, A.; Krummenacker, M.; Latendresse, M.; Midford, P. E. The Ecocyc Database. EcoSal Plus 2018, 8 (1).

(73) Li, G.-W.; Burkhardt, D.; Gross, C.; Weissman, J. S. Quantifying Absolute Protein Synthesis Rates Reveals Principles Underlying Allocation of Cellular Resources. Cell 2014, 157 (3), 624–635.

(74) UniProt: The Universal Protein Knowledgebase in 2021. Nucleic Acids Res 2021, 49 (D1), D480–D489.

(75) Cohen, R. D.; Pielak, G. J. A Cell Is More than the Sum of Its (Dilute) Parts: A Brief History of Quinary Structure. Protein Science 2017, 26 (3), 403–413.

(76) Guin, D.; Gruebele, M. Weak Chemical Interactions That Drive Protein Evolution: Crowding, Sticking, and Quinary Structure in Folding and Function. Chem Rev 2019, 119 (18), 10691–10717.

(77) Gupta, C.; Sarkar, D.; Tieleman, D. P.; Singharoy, A. The Ugly, Bad, and Good Stories of Large-Scale Biomolecular Simulations. Curr Opin Struct Biol 2022, 73, 102338.

(78) Rickard, M. M.; Zhang, Y.; Gruebele, M.; Pogorelov, T. v. In-Cell Protein–Protein Contacts: Transient Interactions in the Crowd. J Phys Chem Lett 2019, 10 (18), 5667– 5673.

(79) Beck, M.; Baumeister, W. Cryo-Electron Tomography: Can It Reveal the Molecular Sociology of Cells in Atomic Detail? Trends Cell Biol 2016, 26 (11), 825–837.

(80) Tegunov, D.; Xue, L.; Dienemann, C.; Cramer, P.; Mahamid, J. Multi-Particle Cryo-EM Refinement with M Visualizes Ribosome-Antibiotic Complex at 3.5 Å in Cells. Nat Methods 2021, 18 (2), 186–193.

## References

(1) Liang, J.; Zhang, L.; Tan, X.; Qi, Y.; Feng, S.; Deng, H.; Yan, Y.; Zheng, J.; Liu, L.; Tian, C. Chemical Synthesis of Diubiquitin-based Photoaffinity Probes for Selectively Profiling Ubiquitin-binding Proteins. Angewandte Chemie 2017, 129 (10), 2788–2792.

(2) Zheng, Q.; Pang, Z.; Liu, J.; Zhou, Y.; Sun, Y.; Yin, Z.; Lou, Z. Photoaffinity Palladium Reagents for Capture of Protein–Protein Interactions. Org Biomol Chem 2019, 17 (26), 6369–6373.

(3) Shigdel, U. K.; Zhang, J.; He, C. Diazirine-Based DNA Photo-Cross-Linking Probes for the Study of Protein–DNA Interactions. Angewandte Chemie 2008, 120 (1), 96–99.

(4) Ran, C.; Pantazopoulos, P.; Medarova, Z.; Moore, A. Synthesis and Testing of Beta-Cell-Specific Streptozotocin-Derived Near-Infrared Imaging Probes. Angewandte Chemie 2007, 119 (47), 9156–9159.

(5) Ran, C.; Pantazopoulos, P.; Medarova, Z.; Moore, A. Synthesis and Testing of Beta-Cell-Specific Streptozotocin-Derived Near-Infrared Imaging Probes. Angewandte Chemie 2007, 119 (47), 9156–9159.

(6) Gerfaud, T.; Martin, C.; Bouquet, K.; Talano, S.; Millois-Barbuis, C.; Musicki, B.; Boiteau, J.-G.; Cardinaud, I. Process Development and Good Manufacturing Practice Production of a Tyrosinase Inhibitor via Titanium-Mediated Coupling between Unprotected Resorcinols and Ketones. Org Process Res Dev 2017, 21 (4), 631–640.

(7) Diamond, R. Real-Space Refinement of the Structure of Hen Egg-White Lysozyme. J Mol Biol 1974, 82 (3), 371–391.

